# The interplay between focus of attention, respiratory phases, and the Heartbeat Evoked Potential

**DOI:** 10.1101/2023.08.13.553126

**Authors:** Andrea Zaccaro, Francesca della Penna, Elena Mussini, Eleonora Parrotta, Mauro Gianni Perrucci, Marcello Costantini, Francesca Ferri

## Abstract

The Heartbeat Evoked Potential (HEP) is an EEG fluctuation that reflects the cortical processing of cardiac signals. HEP amplitude increases during various tasks involving cardiac interoception. Recent research has also indicated that HEP amplitude and cardiac interoceptive accuracy are higher during exhalation compared to inhalation. This difference may be due to the suppression of heartbeat-related sensations during inhalation and the amplification of sensations during exhalation through attentional mechanisms. Despite significant advancements in HEP research, the interactions between the HEP, interoceptive attention, and respiration are still unclear. In this study, we developed a novel experimental paradigm to investigate the relationship between HEP amplitude and respiratory phases during tasks that involve attention to cardiac interoception, non-cardiac interoception (specifically, respiration), and exteroceptive stimuli. The tasks included the Heartbeat Counting Task and the Breath Counting Task as interoceptive tasks, as well as the Cardiac-Tone Counting Task and the Breath-Tone Counting Task as exteroceptive tasks. Results demonstrated significant increases in HEP amplitude during the Heartbeat Counting Task compared to the Cardiac-Tone Counting Task and the Breath Counting Task, mostly observed over fronto-central electrodes in a late time-window. Notably, the amplitude increases during the Heartbeat Counting Task were primarily driven by HEPs recorded during exhalation, while inhalation had minimal impact. These findings align with the predictive coding model of interoceptive perception, suggesting that HEP amplitude reflects a precision-weighting process of prediction errors related to cardiac sensations that is specifically influenced by attention directed toward the heart. Furthermore, our findings emphasize the crucial role of exhalation in this precision-weighting process. These results may have considerable implications for the development of respiratory interventions to fine-tune cardiac interoception.

## 1. Introduction

The Heartbeat Evoked Potential (HEP) is an event-related potential that is time-locked to the cardiac cycle. It is derived by averaging a large number of EEG epochs surrounding simultaneously recorded ECG R-peaks or T-peaks. The HEP was initially described several decades ago in works by Schandry and colleagues (Schandry & Weitkunat, 1986). Since then, the HEP literature has gradually expanded and in recent years it has gained renewed interest within the field of interoception research (Berntson & Khalsa, 2021; Chen et al., 2021). The HEP is primarily recognized as an electrophysiological signature that reflects the cortical processing of cardiac signals (Pollatos & Schandry, 2004). Its physiological correlates are mainly associated with the rhythmic discharge of baroreceptors located in the aortic arch and carotid arteries (Critchley & Harrison, 2013; Garfinkel & Critchley, 2016; Gray et al., 2007; Silvani et al., 2016). Baroreceptors are stretch receptors that respond to both tonic and phasic pressure changes related to cardiac activity. During diastole, they are relatively inactive, whereas during systole they become highly active, providing information about the cardiac state via the vagus nerve, the nucleus of the solitary tract, and the ventromedial posterior thalamic nucleus (Craig, 2003; Critchley & Harrison, 2013; Garfinkel & Critchley, 2016). Cardiac interoceptive information then reaches various brain regions, including the amygdala, insula, anterior and posterior cingulate cortices, and primary somatosensory cortices. Eventually, this information is detected by the EEG recordings, typically over fronto-central areas (Canales-Johnson et al., 2015; Kern et al., 2013; Park & Tallon-Baudry, 2014, for reviews see Coll et al., 2021; Park & Blanke, 2019).

The predictive coding model of interoceptive perception (Barrett & Simmons, 2015; Pezzulo, 2014; Seth, 2013; Seth & Friston, 2016) has recently provided a fascinating interpretation of the HEP. Similar to exteroception, the brain is viewed as an active generator of interoceptive inferences, and interoceptive perception is believed to rely on prior knowledge and expectations. The brain continuously constructs and updates internal models of the body’s physiological state through an iterative process of Bayesian hypothesis testing and revision. The goal is to minimize the mismatch between internal models and actual interoceptive input, achieved through both perceptual inference and active inference mechanisms (Clark, 2013; Friston, 2010). The ultimate objective is to converge toward the most accurate representation of the body state. Prediction errors that arise within this process can serve two purposes: revising prior beliefs by ascending the hierarchical structure and inducing perceptual modifications, or aligning these beliefs with reality by engaging peripheral reflexes. Within this framework, HEPs can be understood as a precision-weighting process of prediction errors associated with heartbeat sensations. Importantly, interoceptive attention can enhance the precision of interoceptive information by amplifying the weighting of the corresponding prediction errors, resulting in increased HEP amplitude (Ainley et al., 2016; Petzschner et al., 2019). Accordingly, studies have shown that when individuals direct their attention toward the heart while performing various cardiac interoceptive tasks, HEP amplitude increases (García-Cordero et al., 2017; Mai et al., 2018; Montoya et al., 1993; Petzschner et al., 2019; Salamone et al., 2018; Schandry & Weitkunat, 1986; Villena-González et al., 2017; Yuan et al., 2007; Zaccaro et al., 2022).

Our recent research has unveiled an intriguing relationship between HEP amplitude, cardiac interoceptive accuracy, and respiratory phases. Specifically, we observed an enhancement of both HEP amplitude and cardiac interoceptive accuracy during exhalation compared to inhalation while performing the Heartbeat Detection (HBD) task (Zaccaro et al., 2022), which involves focusing attention on the heart and pressing a button in synchrony with it (Fittipaldi et al., 2020; Melloni et al., 2013). These findings were interpreted in accordance with the predictive coding model of interoceptive perception (Al et al., 2020, 2021; Allen, Levy, et al., 2022; Allen, Varga, et al., 2022; Grund et al., 2022). According to this model, the brain continuously receives rhythmic and predictable interoceptive signals from the body, effectively suppressing the associated physiological noise (Birn, 2012). During the interoceptive condition of the HBD task, when attention is directed towards the heart, the brain likely receives both heartbeat-related sensations and respiratory interoceptive signals, the latter being mainly sent upstream and consequentially inhibited during inhalations (Noble & Hochman, 2019; Streeter et al., 2012). As a result, heartbeat-related sensations occurring during inhalations may be suppressed along with respiratory-related sensations. On the other hand, the precision of heartbeat-related sensations during exhalations is enhanced by attention, leading to higher HEPs and improved interoceptive performance (Zaccaro et al., 2022).

Despite significant advancements in HEP research, there is still uncertainty regarding the specific relationship between HEP and different types of interoceptive attention, as well as its interaction with the respiratory phases. For instance, HEP augmentation has been associated not only with attention directed towards the heart but also with the attention focused on the bodily self or the self in a broader sense (Babo-Rebelo, Richter, et al., 2016; Babo-Rebelo, Wolpert, et al., 2016; Park et al., 2016, 2018; Sel et al., 2017). However, while previous studies have compared HEP amplitude between cardiac interoceptive tasks and cardiac exteroceptive tasks (Coll et al., 2021), there has been a lack of research comparing HEP amplitude between cardiac interoceptive tasks and *non-cardiac* interoceptive tasks, which involve attending to interoceptive sensations unrelated to the heart. Furthermore, in our previous study that demonstrated respiratory phase-related changes in HEP amplitude (Zaccaro et al., 2022), there were likely confounding factors due to the inherent motor activity involved in the HBD task. This posed challenges for directly comparing the interoceptive and exteroceptive conditions of the HBD task.

In the present study, we have developed a novel experimental paradigm that allowed us to separate the effects of interoceptive attention from the specific physiological system upon which it was focused on. We adopted the Heartbeat Counting Task (HCT) as the cardiac interoceptive task, thereby circumventing potential confounding effects associated with movement-related artifacts present in the HBD task (Zaccaro et al., 2022). Our first analysis compared HEP amplitude between tasks focused on cardiac interoception and exteroception, and tasks targeting *non-cardiac* interoception and exteroception, specifically respiration. If HEP amplitude indeed serves as a neural correlate specific to cardiac interoceptive prediction error, we expected to observe an increase not only during cardiac interoceptive versus exteroceptive attention but also when comparing it to respiratory interoception, where participants needed to concentrate on their breath. No variation in HEP amplitude was expected between the interoceptive and respiratory exteroceptive tasks. Our second analysis investigated higher-level effects of respiration on HEP responses, along with their interaction with attention. We both replicated and expanded upon our previous findings of HEP amplitude changes across respiratory phases using the HCT. Then, we further explored respiratory phase-dependent modulations of HEPs by directly comparing the cardiac tasks with the respiratory tasks. Our rationale was grounded in the assumption that if the increase in HEP amplitude during exhalations, relative to inhalations, reflects heightened attention allocation and enhanced precision-weighting of heartbeat sensations, this augmentation should be exclusive to the cardiac interoceptive task (HCT). Finally, our experimental design allowed us to assess the contribution of HEPs recorded during different respiratory phases across tasks, specifically determining whether the observed increase in HEP amplitude during the cardiac interoceptive task was primarily driven by HEPs recorded during inhalation or exhalation. We predicted a more prominent role of HEPs recorded during exhalation compared to inhalation.

## 2. Results

### 2.1. Overview

After a baseline consisting in a resting-state condition with eyes open, participants underwent four different tasks, involving both interoceptive and exteroceptive attention (“Attention Focus” factor), and focused on both the cardiac and the respiratory systems (“System” factor). In the cardiac interoceptive task, participants had to focus on their heart, count and report the number of perceived heartbeats (Heartbeat Counting Task, HCT; Pollatos & Schandry, 2004; Schandry & Weitkunat, 1986). In the respiratory interoceptive task, participants had to focus on their breath, count and report the number of their breathing cycles (Breath Counting Task, BCT; Herrero et al., 2018). In the cardiac exteroceptive task participants had to count and report the number of beats they heard in an audio recording of a heartbeat (Cardiac-Tone Counting Task, C-TCT; Wiebking et al., 2015), while in the respiratory exteroceptive task, they had to count and report the number of breaths they heard in an audio recording of a breathing person (Breath-Tone Counting Task, B-TCT, adapted from Herrero et al., 2018). EEG, ECG, and respiratory signals were simultaneously recorded throughout all the experiment.

### 2.2. Heartbeat Evoked Potential activity is related to cardiac interoceptive attention

To test the hypothesis that HEP activity is modulated by cardiac interoceptive attention, we performed a 2×2 repeated measures ANOVA mass univariate test, followed by a cluster-based correction, with mean HEP amplitude as the dependent variable, and System (cardiac vs. respiratory) and Attention Focus (interoceptive vs. exteroceptive) as within-participants factors. The number of included epochs for each task is showed in Table S1. We performed the test over an early time-window and a late time-window (see Section 4.11). In the early time-window (i.e., 150-349 msec after the R-peak), the ANOVA showed a significant System by Attention Focus interaction over Fp1, AF3, F3, F1, FC3, FC1, AFz, Fz, FCz, AF4, and F2 (F_1,31_ = 6.906, p_clust_ = .008, *η*2p = .182). There was no significant main effect of System (F_1,31_ = .841, p_clust_ = .366, *η*2p = .026) and Attention Focus (F_1,31_ = .039, p_clust_ = .844, *η*2p = .001). Despite the significant interaction, planned t-tests revealed that there were no significant differences between the following conditions: HCT vs. C-TCT (t_31_ = -.582, p_clust_ = .565, Cohen’s d = .103), HCT vs. BCT (t_31_ = −2.109, p_clust_ = .087, Cohen’s d = .373), BCT vs. B-TCT (t_31_ = .965, p_clust_ = .342, Cohen’s d = .171), and C-TCT vs. B-TCT (t_31_ = 1.113, p_clust_ = .274, Cohen’s d = .197).

For the late time-window, the mean HEP amplitude between 350 and 600 msec after the R-peak was set as the dependent variable. The ANOVA did not reveal a main effect of System (F_1,31_ = .573, p_clust_ = .266, *η*2p = .018) or Attention Focus (F_1,31_ = 3.264, p_clust_ = .081, *η*2p = .095). However, the System by Attention Focus interaction was significant (F_1,31_ = 10.09, p_clust_ = .006, *η*2p = .246, Fig. 1A) over the channel Fp1, AFz, AF3, AF4, Fz, F1, F2, F3, FCz, FC1, and FC3. Crucially, planned t-tests revealed significant increases in HEP activity during the HCT compared to the C-TCT over the F3, Fz, AFz, F1, F2, AF3, AF4, Fpz, and FCz channels (t_31_ = 2.761, p_clust_ = .006, Cohen’s d = .488, Fig. 1B, 1F), as well as compared to the BCT over the channel F3, Fz, AFz, F1, F2, AF3, and AF4 (t_31_ = 2.32, p_clust_ = .028, Cohen’s d = .41, Fig. 1C). No significant differences were observed between the BCT vs. B-TCT (t_31_ = .627, p_clust_ = .535, Cohen’s d = .111, Fig. 1D), nor between the C-TCT vs. B-TCT (t_31_ = −1.33, p_clust_ = .215, Cohen’s d = .235, Fig. 1E). These results indicate that the cardiac interoceptive task (HCT) induced significant HEP increases compared to both the cardiac exteroceptive (C-TCT) and respiratory interoceptive (BCT) tasks in the late time-window (i.e., 350-600 msec after the R-peak). For this reason, we performed further analyses focusing on the late time-window.

**Figure 1.**
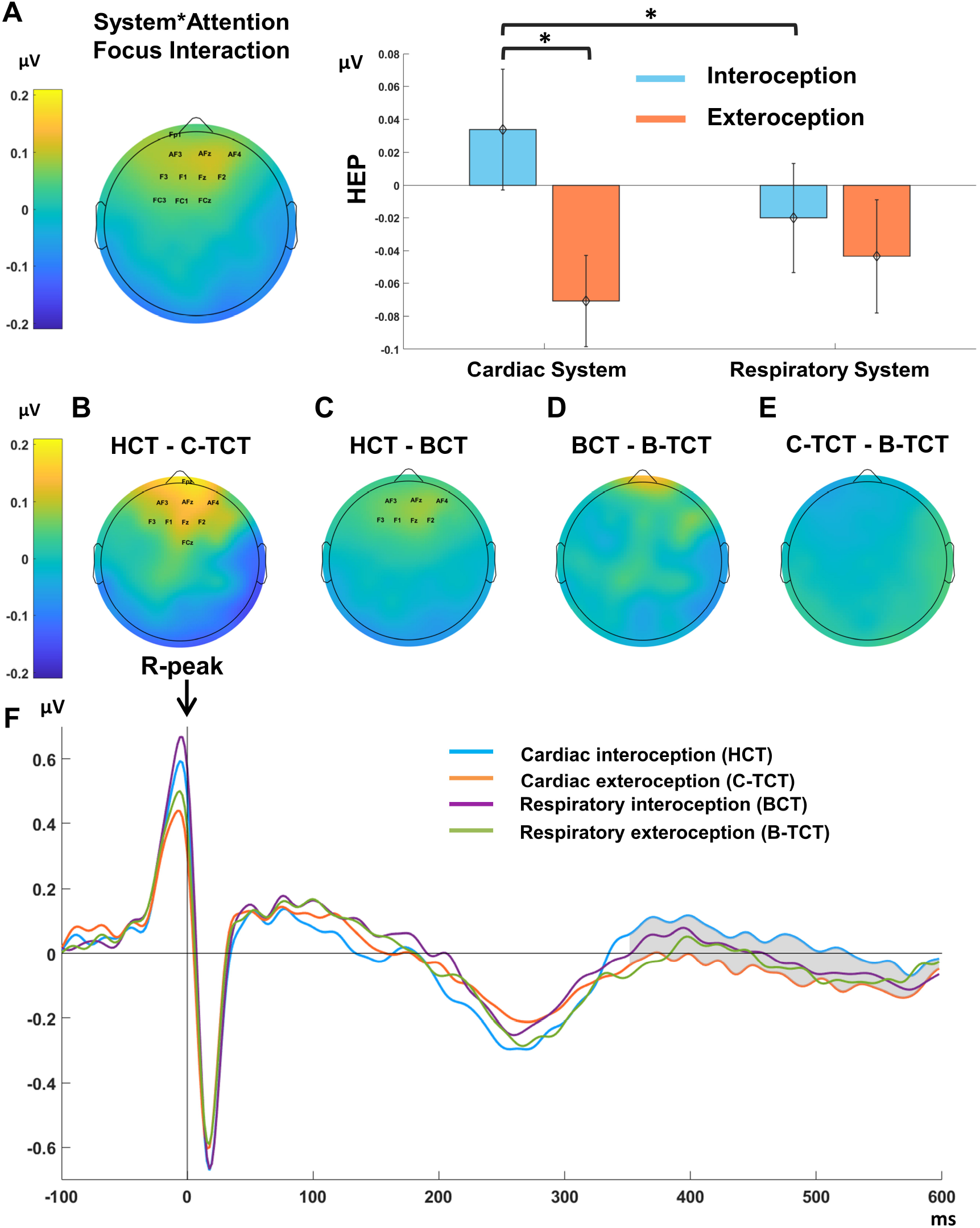
HEP activity is related to cardiac interoceptive attention. **(A-left)** Topographical scalp distribution showing the significant interaction between System*Attention Focus factors. Mean HEP amplitude in the significant time window (350-600 msec after the R-peak) is calculated with the following formula: (Cardiac Interoception – Cardiac Exteroception) – (Respiratory Interoception – Respiratory Exteroception). Significant interaction was detected over highlighted electrodes (Fp1, AF3, AFz, AF4, F3, F1, Fz, F2, FC3, FC1, and FCz). **(A-right)** Mean HEP amplitude assessed over significant electrodes across the four tasks. Significant planned t-tests are indicated by asterisks (* p < .05). **(B)** Topographical scalp distribution showing mean HEP differences (350 - 600 msec after the R-peak) between the HCT vs. C-TCT (significant differences were detected over highlighted electrodes: Fpz, AF3, AFz, AF4, F3, F1, Fz, F2, and FCz), **(C)** HCT vs. BCT (significant differences were detected over highlighted electrodes: AF3, AFz, AF4, F3, F1, Fz, and F2), **(D)** BCT vs. B-TCT, and **(E)** C-TCT vs. B-TCT. **(F)** Grand-average HEP waveforms pooled for significant electrodes. The time courses of the HEP are shown for the HCT (light blue), C-TCT (orange), BCT (purple), and B-TCT (green). The grey area marks the time window of significant differences in the contrast between the HCT and C-TCT. Abbreviations: HCT-Heartbeat Counting Task, C-TCT-Cardiac-Tone Counting Task, BCT-Breath Counting Task, B-TCT-Breath-Tone Counting Task.

### 2.3. Heartbeat Evoked Potential activity is related to the respiratory phase

We tested whether HEP activity in the late time-window was associated with the respiratory phase during the performance of the four tasks. To this aim, we differentiated HEPs occurring during inhalation and exhalation phases. We compared the number of analysed HEP epochs between these phases within the baseline and each of the four tasks. In general, we found more HEP epochs during inhalation than exhalation (Table S2). However, we focused on mean HEP amplitude rather than peak amplitude, as it is less influenced by differences in the number of trials (Luck, 2014). We firstly replicated our previous finding (Zaccaro et al., 2022) by running a cluster-based permutation t-test between 350 and 600 milliseconds following the R-peak at baseline. This analysis showed higher HEP amplitude during the exhalation phase than the inhalation phase. Specifically, we found higher HEP activity in a cluster of central, parietal, and parieto-occipital electrodes (C2, CPz, CP1, CP4, Pz, P1, P2, P3, P4, POz, PO3, and PO4) (t_27_ = 3.33, p_clust_ = .005, Cohen’s d = .629, Fig. S1).

Then, we conducted a 2×2×2 mass univariate repeated measures ANOVA followed by cluster-based correction with mean HEP amplitude as the dependent variable, and System (cardiac vs. respiratory), Attention Focus (interoceptive vs. exteroceptive), and Phase (inhalation vs. exhalation) as within-subject factors. This analysis focused on the significant time interval (late time-window) and channels (fronto-central regions) identified in the previous analysis, which revealed interoceptive attention-driven increases in HEP activity. As expected, the results indicated no main effect of System (F_1,27_ = 2.466, p_clust_ = .128, *η*2p = .084) or Attention Focus (F_1,27_ = 1.459, p_clust_ = .237, *η*2p = .051), but a significant interaction between System and Attention Focus (F_1,27_ = 5.788, p_clust_ = .01, *η*2p = .177) over Fp1, AF3, F5, F3, F1, AFz, Fz, AF4, and F2. These trivial findings mirrored the previous analysis that did not consider respiratory phases. In addition, no main effect of Phase (F_1,27_ = .102, p_clust_ = .752, *η*2p = .004), interaction effects between Attention Focus and Phase (F_1,27_ = 1.563, p_clust_ = .222, *η*2p = .055), or between System and Phase (F_1,27_ = 1.067, p_clust_ = .311, *η*2p = .038) were observed. Crucially, the three-way interaction of System by Attention Focus by Phase was significant over the channels Fp2, AFz, AF3, AF4, Fz, and F4 (F_1,27_ = 6.771, p_clust_ = .043, *η*2p = .2, Fig. 2A). Follow-up planned t-tests confirmed that HEP activity during exhalation, compared to inhalation, was significantly increased during the cardiac interoceptive task (HCT) over Fpz, Fp2, AFz, F2, and FC2 (t_27_ = 2.199, p_clust_ = .037, Cohen’s d = .42, Fig. 2B, Fig. 2G). Conversely, there were no significant changes of HEP activity between exhalation and inhalation during the respiratory interoceptive task (BCT) (t_27_ = .742, p_clust_ = .464, Cohen’s d = .14), the cardiac exteroceptive task (C-TCT) (t_27_ = .86, p_clust_ = .397, Cohen’s d = .163), or the respiratory exteroceptive task (B-TCT) (t_27_ = .006, p_clust_ = .284, Cohen’s d = .001). Interestingly, HEPs recorded during exhalation in the HCT were significantly higher than those recorded during exhalation in the C-TCT (t_27_ = 2.24, p_clust_ = .04, Cohen’s d = .423, Fig. 2C) over Fpz, Fp2, AFz, AF4, and Fz, as well as higher than those recorded during exhalation in the BCT (t_27_ = 3.476, p_clust_ = .005, Cohen’s d = .657, Fig. 2D) over Fp2, AFz, AF3, AF4, Fz, F1, F2, F3, and F4. Finally, there were no significant changes in HEP activity when comparing HEPs recorded during inhalation in the HCT with those recorded during inhalation in the C-TCT (t_27_ = .743, p_clust_ = .464, Cohen’s d = .14, Fig. 2E) or the BCT (t_27_ = .19, p_clust_ = .85, Cohen’s d = .036, Fig. 2F). These results replicate our previous study (Zaccaro et al., 2022) demonstrating higher HEP activity during exhalation compared to inhalation in a cardiac interoceptive task (HCT). Importantly, they further indicate that this respiratory phase-dependent modulation of HEP activity does not extend to the respiratory interoceptive task (BCT). Crucially, these findings reveal that the previously observed HEP amplitude increases during the HCT compared to the C-TCT and the BCT are primarily due to HEPs recorded during the exhalation phase of respiration, with negligible contributions of HEPs recorded during inhalation.

**Figure 2.**
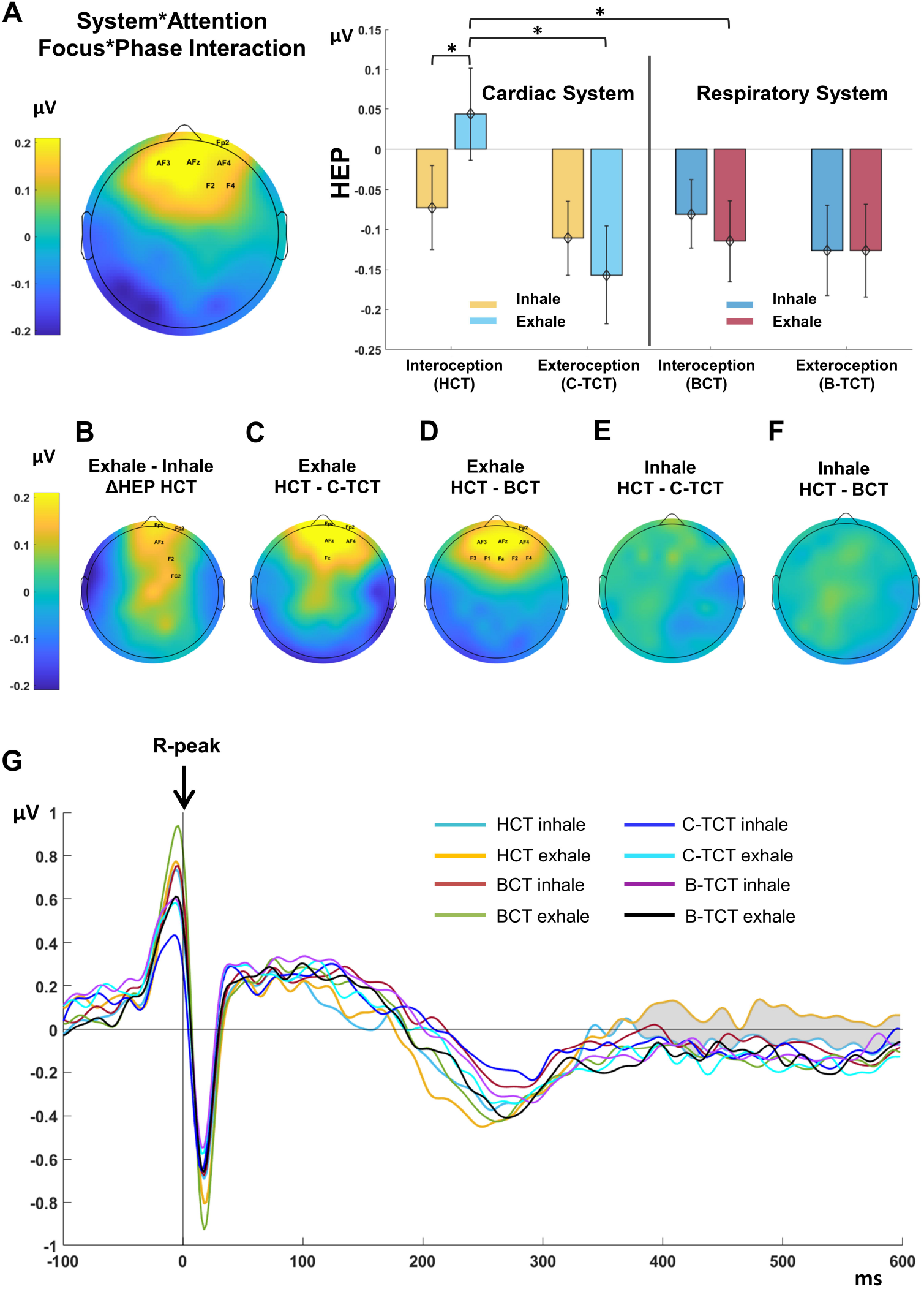
HEP activity is related to the respiratory phase during the HCT. **(A-left)** Topographical scalp distribution showing the significant interaction between System*Attention Focus*Phase factors. Mean HEP amplitude in the significant time window (350-600 msec after the R-peak) is calculated with the following formula: [(Cardiac Interoception Exhale – Cardiac Interoception Inhale) – (Cardiac Exteroception Exhale – Cardiac Exteroception Inhale)] - [(Respiratory Interoception Exhale – Respiratory Interoception Inhale) – (Respiratory Exteroception Exhale – Respiratory Exteroception Inhale)]. Significant interaction was detected over highlighted electrodes (Fp2, AF3, AFz, AF4, F2, and F4). **(A-right)** Mean HEP amplitude assessed over significant electrodes across the four tasks and respiratory phases. Significant planned t-tests are indicated by asterisks (* p < .05). **(B)** Topographical scalp distribution showing mean HEP differences (350 – 600 msec after the R-peak) between exhale and inhale phase (ΔHEP) during the HCT (significant differences were detected over highlighted electrodes: Fpz, Fp2, AFz, F2, and FC2). **(C)** Topographical scalp distribution showing mean HEP differences (350 – 600 msec after the R-peak), during exhalation between the HCT and C-TCT (significant differences were detected over highlighted electrodes: Fpz, Fp2, AFz, AF4, and Fz), **(D)** during exhalation between the HCT and BCT (significant differences were detected over highlighted electrodes: Fp2, AF3, AFz, AF4, F3, F1, Fz, F2, and F4), **(E)** during inhalation between the HCT and C-TCT, and **(F)** during inhalation between the HCT and BCT. **(G)** Grand-average HEP waveforms pooled for significant electrodes. The time courses of the HEP are shown for the HCT-inhale (turquoise), HCT-exhale (yellow), BCT-inhale (dark red), BCT-exhale (green), C-TCT-inhale (blue), C-TCT-exhale (light blue), B-TCT-inhale (purple), B-TCT-exhale (black). The grey area marks the time window of significant differences in the contrast between exhale and inhale phase (ΔHEP) during the HCT. Abbreviations: HCT-Heartbeat Counting Task, C-TCT-Cardiac-Tone Counting Task, BCT-Breath Counting Task, B-TCT-Breath-Tone Counting Task.

### 2.4. Heartbeat Evoked Potentials and cardio-respiratory physiology

Potential factors that modulate HEP activity and/or cardiac interoception are represented by the heart stroke volume (indexed by the ECG signal amplitude and the HR) and the respiratory rate (Buot et al., 2021; Candia-Rivera et al., 2022; Coll et al., 2021; Dirlich et al., 1997; Kern et al., 2013; MacKinnon et al., 2013; Smith et al., 2020, 2021; Leganes-Fonteneau et al., 2021). To assess whether observed changes in HEP amplitude were related to concurrent modifications in heart activity and physiology, we performed control analyses comparing participants’ ECG signal amplitude as well as cardiac features of interest across System and Attention Focus levels (Table S3). Time-domain parameters, such as Heart Rate (HR), and frequency-domain parameters, such as power in High Frequency band log value (HFlog), Heart Rate Variability (HRV) total power, and Low Frequency/High Frequency ratio (LF/HF) were computed. Respiratory features of interest were the breathing rate, the average inhale and exhale duration, and the Inhalation/Exhalation (I/E) ratio (Supplementary Material 1). ECG amplitude was unchanged across the four tasks (Fig. S2), while the mean HR was specifically lower during the HCT, compared to all the other tasks (Fig. S3A). The other cardio-respiratory features showed a non-specific main effect for the interoceptive vs. exteroceptive tasks (Supplementary Material 2, Fig. S3). Given its specific reduction during the HCT, we controlled for instantaneous HR influences on HEPs at the single-trial level by running a linear mixed-effects model analysis comprising 91980 trials (each trial corresponds to a HEP epoch). The analysis showed a negligible effect of instantaneous HR on HEP activity across the four tasks (Supplementary Material 3, Table S4, Fig. S4). Among respiratory phases, we performed additional analyses to control for confounding influence of cardiac physiology on HEP activity, as indexed by phase-specific HR and ECG amplitude (Buot et al., 2021), to support the assumption that observed differences in HEP amplitude between inhalation and exhalation were not driven by heart stroke volume modifications (Supplementary Material 4). At the subject level, we found that the ECG signal amplitude was higher during the exhalation phase, compared to inhalation, across the four tasks (Fig. S5, Fig. S6A). When considering the HR, we found a specific reduction at exhalation during the HCT compared to the C-TCT and the BCT (Fig. S6B). Therefore, at the single-trial level, we controlled for cardiac influences on HEPs across the respiratory cycle (inhalation vs. exhalation), as assessed with instantaneous HR and trial-based ECG amplitude, by running a linear mixed-effects model analysis comprising 17196 trials during the performance of the HCT (Supplementary Material 5, Table S5, Fig. S7). Results showed no interaction between HR, ECG amplitude, and respiratory phases, suggesting that these covariates do not mediate the effect of the respiratory phase on HEP amplitude during the HCT.

### 2.5. Respiratory cycle analyses during the Heartbeat Counting Task

To capture the smooth changes in HEP amplitude across the respiratory cycle, we employed circular statistics and divided the respiratory cycle into phase bins (Berens, 2009; Park et al., 2020). We focused on the cardiac interoceptive task (HCT) due to the significant HEP changes observed among respiratory phases. Following the circular analysis approach, we characterized HEP activity over the entire respiratory cycle, from one inhalation to the next one. Then, we used a median split method to split HEP values into two bins of high-amplitude HEPs and low-amplitude HEPs, for each participant (Al et al., 2020, 2021). Subsequently, we calculated the mean angle direction for low-amplitude HEPs and high-amplitude HEPs for each participant. To test whether the distribution of high and low HEPs deviated from the uniform distribution across the respiratory cycle at the group level, we used the Hodges-Ajne test (or Omnibus test, Park et al., 2020). The omnibus test showed that uniformity was rejected for both low- and high-amplitude HEPs (low-amplitude HEPs: M = 1, p < .001; high-amplitude HEPs: M = 6, p = .044). Consistent with the binary analysis results, the distribution of low-amplitude HEPs was clustered around the middle phase of inhalation (first and second quadrant; mean angle = 96.33°, Fig. 3A). Conversely, the distribution of high-amplitude HEPs was clustered around the early phase of inhalation (first quadrant) and extended to the exhalation phase (third and fourth quadrant), with the mean direction occurring near the transition from exhalation to inhalation (mean angle = 22.98°, Fig. 3B). Confirming these results, the Watson-Williams test revealed significant difference between low- and high-amplitude HEPs (Watson-William’s test: F_1,54_ = 11.061, p = .002).

**Figure 3.**
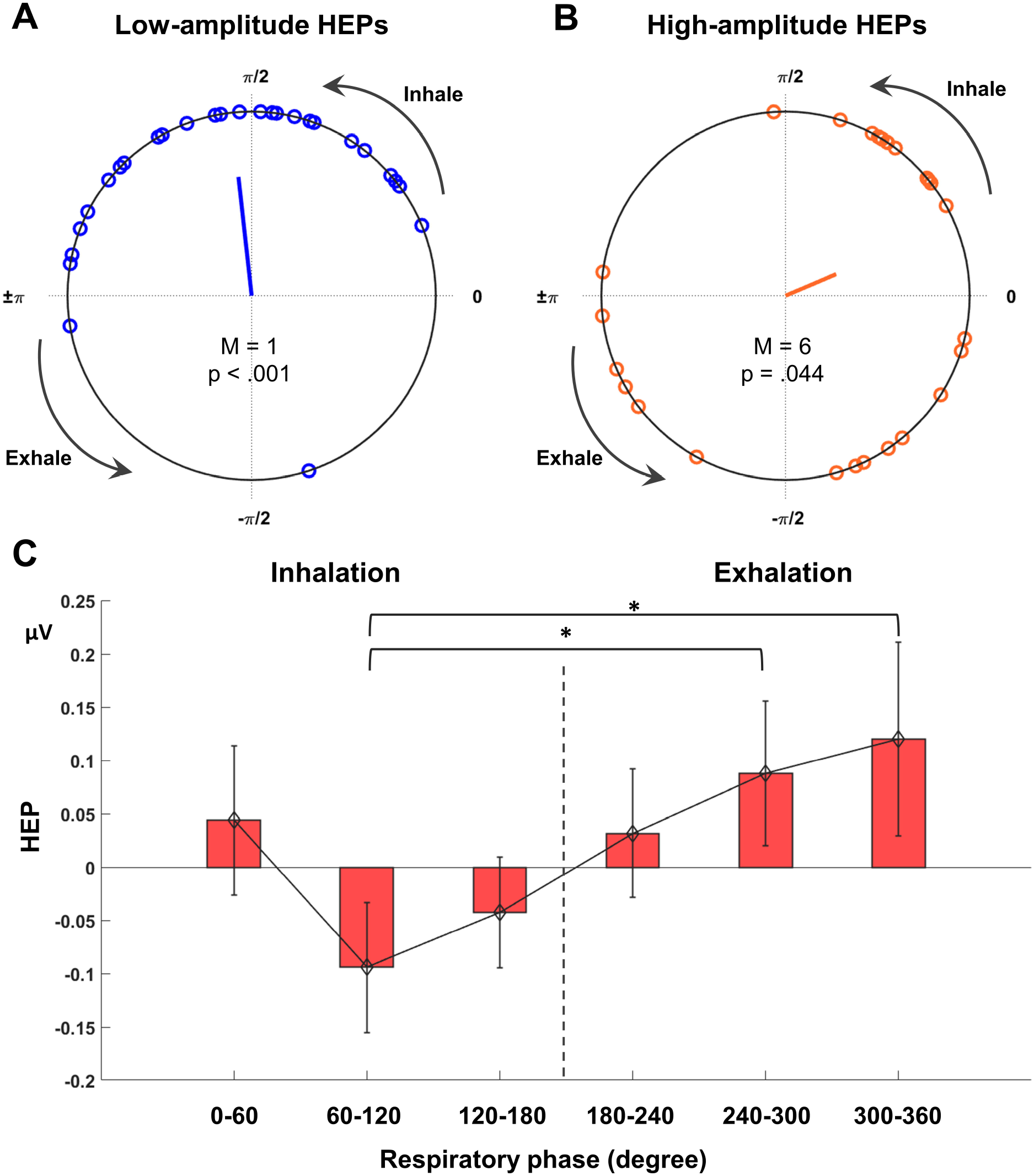
Respiratory cycle analyses. Circular distribution of mean angles (HEPs onsets relative to the respiratory cycle) for **(A)** low-amplitude HEPs (blue), and **(B)** high-amplitude HEPs (orange). Zero degree corresponds to inhalation onset. Each dot indicates the mean angle of one participant. The direction of the line in the centre indicates the mean angle across the participants while the arrow length represents the mean resultant length. The resulting M value and p value of the Omnibus test are indicated below. **(C)** The mean amplitude of HEP as a function of six equally sized bins of the respiratory phase. Zero degree corresponds to inhalation onset. Error bars represent the standard error. Significant planned t-tests are indicated by asterisks (* p < .05).

We then assessed the effects of respiratory phases on HEP amplitude during the HCT by analysing HEPs within six equally-sized respiration phase bins covering the 0°-360° interval (0°-60°, 60°-120°, and 120°-180° for inhalation, and 180°-240°, 240°-300°, and 300°-360° for exhalation). Consistent with the binary analysis, a one-way repeated measures ANOVA showed significant changes in HEP amplitude across the respiratory cycle during the HCT (F_5,135_ = 2.45, p = .037, *η*2p = .083). Planned t-tests comparisons between inhalation phase bins and exhalation phase bins confirmed that HEPs detected in the 60°-120° range (middle inhalation) had significantly lower amplitude than HEPs detected in the 240°-300° range (middle exhalation) (t_27_ = −2.713, p = .006, p_FDR_ = .045, Cohen’s d = .513) and in the 300°-360° range (t_27_ = −2.488, p = .01, p_FRD_ = .045, Cohen’s d = .47) (late exhalation) (Fig. 3C). All the other comparisons did not show significant differences after FDR correction (0°-60° vs. 180°-240°: t_27_ = .164, p = .565, Cohen’s d = .031; 0°-60° vs. 240°-300°: t_27_ = −.656, p = .259, Cohen’s d = .124; 0°-60° vs. 300°-360°: t_27_ = −1.043, p = .153, Cohen’s d = .197; 60°-120° vs. 180°-240°: t_27_ = −1.836, p = .039, p_FDR_ = .085, Cohen’s d = .347; 120°-180° vs. 180°-240°: t_27_ = −1.126, p = .135, Cohen’s d = .213; 120°-180° vs. 240°-300°: t_27_ = −1.733, p = .047, p_FDR_ = .085, Cohen’s d = .327; 120°-180° vs. 300°-360°: t_27_ = −2.006, p = .027, p_FDR_ = .081, Cohen’s d = −.379).

### 2.6. Correlation with behavioural and psychometric data

Task accuracy was influenced by System and Attention Focus (Table S6). We found no relationship between HEP activity and task accuracy or confidence (Supplementary Material 6). Given the previous associations of HEP activity with specific self-reported interoceptive sensibility and mindfulness measures (Baranauskas et al., 2017; Billeci et al., 2021; Verdonk et al., 2021), we examined the correlations between self-reported scores obtained on the Trusting and Not-Worrying scales of the MAIA and the total score on the FFMQ, with HEP amplitude. In particular, we investigated the correlations between each of the above-mentioned scales and the mean HEP amplitude recorded during inhalation and exhalation while performing the HCT. Twenty-five participants completed self-report questionnaires. HEP values computed during inhalation in the HCT did not correlate with any of the scales (FFMQ Total Score Pearson’s r = .007, p = .973, Fig. 4A; MAIA Trusting Pearson’s r = .057, p = .787, Fig 4B; MAIA Not-Worrying Pearson’s r = −.357, p = .079, Fig. 4C). In contrast, HEP values computed during exhalation exhibited a positive correlation with the FFMQ total score (Pearson’s r = .493, p = .012, p_FDR_ = .036, Fig. 4D) and the MAIA Trusting scale (Pearson’s r = .447, p = .025, p_FRD_ = .038, Fig. 4E), but not with the MAIA Not-Worrying scale (Pearson’s r = .142, p = .499, Fig. 4F). These findings demonstrate that the relationship between HEP amplitude, interoceptive sensibility, and mindfulness level is specific to HEPs recorded during exhalation, while no significant relationship was observed for HEPs recorded during inhalation.

**Figure 4.**
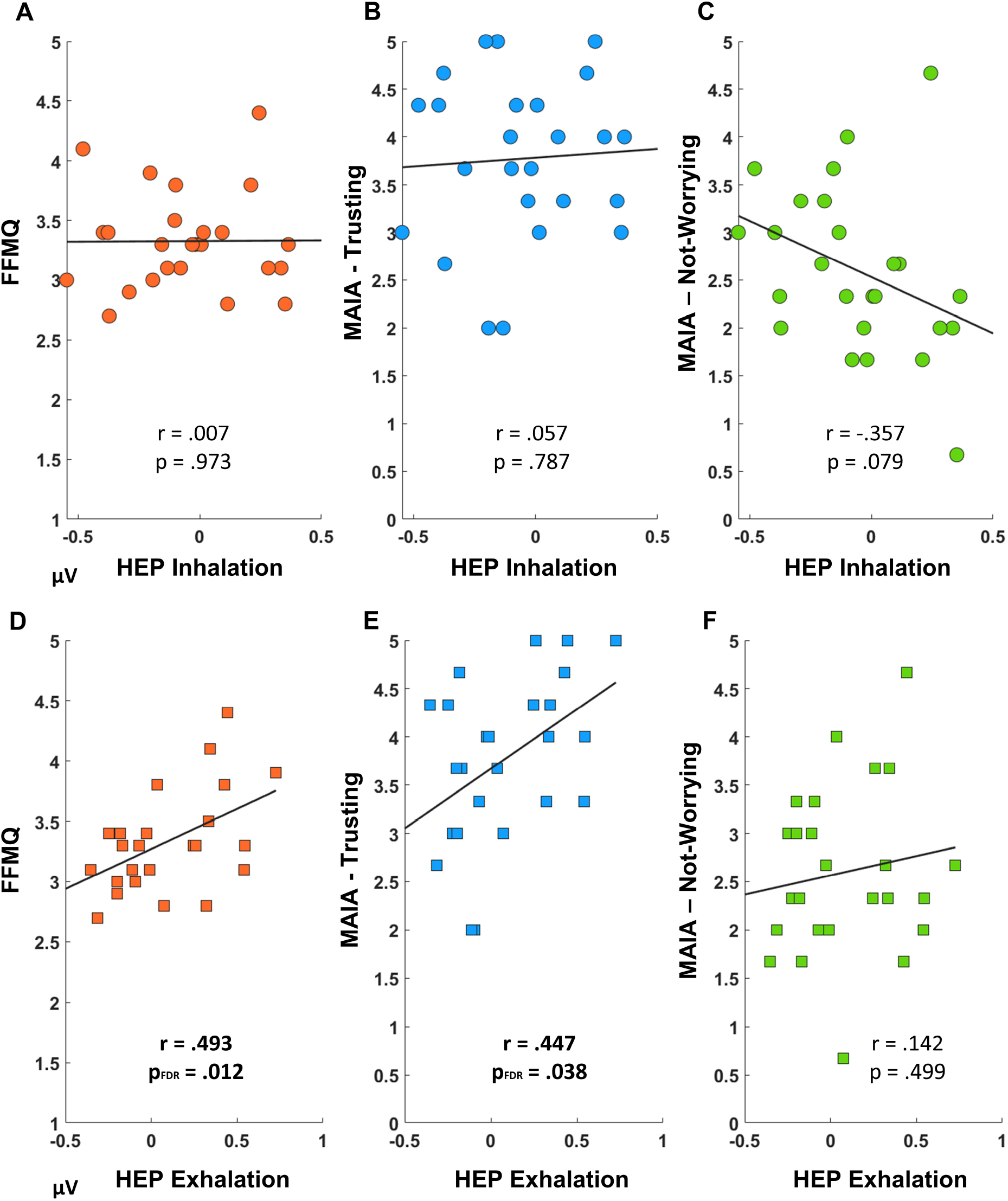
Correlation between psychometric questionnaires and HEPs recorded at inhalation and exhalation. Scatter plots of the linear relationships (Pearson’s correlation) between mean HEP amplitude recorded during inhalation and exhalation and the FFMQ total score, MAIA Trusting scale, and MAIA Not-Worrying scale.

## 3. Discussion

The current study employed a novel experimental design to assess Heartbeat Evoked Potential (HEP) activity in healthy volunteers across four tasks designed to elicit either interoceptive or exteroceptive attention. The study focused on either the cardiac or the respiratory system. The interoceptive tasks included the Heartbeat Counting Task (HCT; Pollatos & Schandry, 2004; Schandry & Weitkunat, 1986) and the Breath Counting Task (BCT; Herrero et al., 2018), while the exteroceptive tasks consisted of the Cardiac-Tone Counting Task (C-TCT; Wiebking et al., 2015) and the Breath-Tone Counting Task (B-TCT; adapted from Herrero et al., 2018). The study aimed to achieve two main objectives. Firstly, it sought to investigate the specificity of HEP augmentation concerning cardiac interoceptive attention. Secondly, it aimed to characterize modulations of HEP amplitude related to respiratory phases during different interoceptive and exteroceptive attentional tasks.

The results of the study revealed significant increases in HEP amplitude during the cardiac interoceptive task (HCT), compared to both the cardiac exteroceptive task (C-TCT) and the respiratory interoceptive task (BCT). These increases of HEP were observed over fronto-central electrodes and were detected in the late time-window, ranging from 350 to 600 milliseconds after the R-peak. In contrast, directing attention towards breathing sensations during the BCT did not elicit significant modulation of HEP amplitude. These findings align with previous studies on HEPs, which also reported attention-related modulation of HEP amplitude during a cardiac interoceptive task in the late time-windows and over fronto-central electrode sites (Baranauskas et al., 2017; Gray et al., 2007; Kumral et al., 2022; Petzschner et al., 2019; Pollatos et al., 2016; Schandry & Montoya, 1996; Schulz et al., 2018; Simor et al., 2021). This supports the hypothesis that HEPs are associated with a precision-weighting process of prediction errors related to cardiac sensations and suggests that this precision-weighting process of prediction errors is enhanced by top-down attention directed towards the heart (Ainley et al., 2016; Petzschner et al., 2019). Furthermore, these results significantly contribute to the current understanding of HEPs by demonstrating that augmentation of HEP amplitude is specifically associated with the allocation of attention to cardiac sensations, as opposed to other forms of interoceptive attention, such as respiratory interoception. Additionally, our data suggest that despite certain limitations that may affect its validity in assessing cardiac interoceptive accuracy (Brener & Ring, 2016; Desmedt et al., 2018; Zamariola et al., 2018), the HCT is a suitable task for eliciting cardiac interoceptive attention.

After selectively assessing HEP activity across tasks, we directly compared respiratory phase-induced modulations of HEPs between the HCT, the BCT, the C-TCT, and the B-TCT. The findings revealed greater respiratory phase-related changes in HEPs (ΔHEP) during the cardiac interoceptive task (HCT) over frontal electrodes. Specifically, during the HCT, the HEP exhibited significantly higher amplitudes during exhalation compared to inhalation in the late time-window (i.e., 350-600 msec) over fronto-central areas. In contrast, no significant changes in HEP activity were observed between exhalation and inhalation during the BCT, the C-TCT, and the B-TCT. Additional analyses using circular statistics (Berens, 2009) supported results obtained from the binary analysis (inhalation vs. exhalation comparison). Specifically, lower-amplitude HEPs were clustered around the central part of the inhalation phase, while higher-amplitude HEPs were concentrated around the exhalation phase and the transition from exhalation to inhalation. By dividing each breath into six equal segments spanning the full 0°-360° range, we observed a gradual increase in HEP amplitude across the respiratory cycle, peaking at the end of the exhalation phase (the 300°-360° segment). Importantly, our experimental design allowed for a direct comparison of HEP amplitude occurring between the inhalation and exhalation phases of respiration during the four tasks. Consequently, we found that HEPs recorded during exhalation in the cardiac interoceptive task (HCT) exhibited significantly higher amplitude than HEPs recorded during exhalation in both the cardiac exteroceptive task (C-TCT) and the respiratory interoceptive task (BCT). Notably, no differences were found when comparing HEPs recorded during inhalation in the HCT with HEPs recorded during inhalation in the C-TCT and the BCT. These results not only replicate but also build upon our previous findings concerning changes in HEP amplitude across respiratory phases (Zaccaro et al., 2022). Firstly, they corroborate previous results by using a different interoceptive task, specifically the HCT, in addition to the previously used HBD task. Secondly, they establish that the modulations of HEP activity based on the respiratory phase are specific to the *cardiac* interoceptive task and do not extend to *non-cardiac* interoceptive tasks, such as the BCT. Lastly, they indicate that, during the cardiac interoceptive task, the brain does not enhance the precision of heartbeat-related sensations during inhalations; rather, it enhances the precision of heartbeat-related sensations exclusively during exhalation.

An additional noteworthy finding is the positive association observed between HEP activity and specific well-being indicators, namely interoceptive sensibility and dispositional mindfulness (Baranauskas et al., 2017; Billeci et al., 2021; Verdonk et al., 2021). During the performance of the cardiac interoceptive task (HCT), we found that HEP amplitude recorded during exhalation was specifically related to self-reported scores at the Trusting and Not-Worrying scales of the Multidimensional Assessment of Interoceptive Awareness (MAIA; Mehling et al., 2012), as well as to the total score of the Five Facet Mindfulness Questionnaire (FFMQ; Baer et al., 2006). In contrast, we did not find significant associations between HEP amplitude during inhalation and these indices. This suggests that HEPs assessed during exhalation are more strongly connected to interoceptive sensibility and mindfulness, which are crucial indicators of well-being and mental health (Bonaz et al., 2021; Hölzel et al., 2011; Khalsa et al., 2018; Paulus, 2013; Tang et al., 2015; Tsakiris & Critchley, 2016). These results could help reconcile conflicting findings in current research on interoceptive improvements in mindfulness practitioners, as some studies have reported non-significant increases in cardiac interoceptive accuracy (see Khalsa et al., 2020, and Treves et al., 2019 for meta-analyses). The lack of significant findings in those studies might be attributed to the failure to consider respiratory phases. Moreover, these findings may provide an explanation for previous studies that did not find associations between trait mindfulness and overall HEP activity (Verdonk et al., 2021). It is plausible that cardiac interoception in meditators is specifically enhanced during exhalation in the long term. However, existing studies have consistently assessed cardiac interoception without distinguishing between respiratory phases, making it challenging to draw definitive conclusions about the enhancement (or lack thereof) of HEP activity during mindfulness practice. Future studies may shed light on this matter.

Taken together, our findings represent a significant advancement in understanding the physiological mechanisms underlying the HEP. Specifically, we can conclude that the overall increases in HEP amplitude observed during the cardiac interoceptive task (HCT), compared to the cardiac exteroceptive task (C-TCT) and the respiratory interoceptive task (BCT), are primarily driven by HEPs recorded during the exhalation phase of respiration, with minimal contribution from HEPs recorded during inhalation. This strengthens our previous interpretation of the variation in interoception across the respiratory cycle during a cardiac interoceptive task (Zaccaro et al., 2022), and introduces a new perspective on HEP activity that emphasizes the crucial role of exhalation. Our data demonstrate that HEP amplitude increases occur only when attention is focused on the heart, specifically during exhalation. This is consistent with the interoceptive predictive processing framework, suggesting that HEP amplitude reflects a precision-weighting process of prediction errors, which is influenced by attention directed toward cardiac sensations. Exhalation seems to enable this precision-weighting mechanism. In other words, exhalation may serve as more than just a respiratory phase that enhances attention towards cardiac interoception; it is a necessary condition for the occurrence of the precision-weighting mechanism of prediction errors related to cardiac sensations. If the first scenario were true, during the HCT task, we would expect to observe lower HEP amplitude during inhalation compared to exhalation, although still higher than the HEP amplitude recorded during inhalation in the other tasks (C-TCT and BCT), considering the attention directed towards the heart. However, we did not observe any differences in HEP amplitude during inhalation across the tasks. Several factors could account for this phenomenon. One higher-level explanation is in line with a recent hypothesis regarding the role of respiration in interoceptive perception (Molle & Coste, 2022). According to this hypothesis, the brain constantly engages in a trade-off between directing attention to interoceptive or exteroceptive stimuli. Inhalation may prioritize external perception by redirecting attention to the surrounding environment, while exhalation may optimize interoception by allocating attentional resources to bodily sensations. Consequently, the imbalance in attentional resource allocation towards exteroceptive functions during inhalation (e.g., visual, Kluger et al., 2021; somatosensory, Grund et al., 2022; and auditory processing, Johannknecht & Kayser, 2022) may disrupt the processing of interoceptive signals, especially in the case of cardiac sensations, which are known to be relatively ambiguous. This effect likely persists regardless of the specific attentional focus on the heart. This explanation aligns with the observed localization of respiratory phase-induced effects on the HEP, particularly over frontal and prefrontal electrodes, which coincide with the Anterior Cingulate Cortex (ACC). The ACC is a well-established cortical source of the HEP (Babo-Rebelo, Richter, et al., 2016; Park et al., 2014, 2016), and has been associated with the precision-weighting of inferred interoceptive states through neuromodulatory gain control, particularly involving noradrenaline and dopamine modulation (Allen, Levy, et al., 2022; Fardo et al., 2017; Feldman & Friston, 2010). Additionally, the ACC is involved in higher-level functions like voluntary top-down attention (Allen et al., 2017; Fleming & Dolan, 2012). Alternatively, the mechanism could be more peripheral and related to the strong connection between the respiratory cycle and the baroreflex, known as Respiratory Sinus Arrhythmia (RSA; Brecher & Hubay, 1955). RSA leads to faster, but inherently weaker heartbeats during inhalation compared to exhalation (Larsson et al., 2021), which may result in reduced HEP amplitude during inhalation. However, the observation that heartbeats vary in strength among respiratory phases does not exclude the possibility of top-down attention being similarly modulated. Instead, it aligns with an adaptive embodied mechanism that optimizes attention in synchrony with respiration. Accordingly, during inhalation, when cardiac interoceptive signals are diminished, attention may be more readily directed towards the external environment, also due to reduced interference from interoceptive signals. Conversely, during exhalation, when cardiac signals are stronger, attention may be more easily directed towards internal processes, thereby enhancing the processing of already heightened interoceptive information. This is consistent with recent studies that have observed positive effects of exhalation on *non-exteroceptive* brain functions, such as associative conditioned learning (Waselius et al., 2019, 2022), voluntary actions (Park et al., 2020), and mental imagery of voluntary actions (Park et al., 2022).

Regardless of the specific mechanism involved, it is crucial for future studies to differentiate between HEPs recorded during different respiratory phases, as exhalation has consistently demonstrated stronger and more specific associations with interoceptive attention and sensibility compared to inhalation. Additionally, it is important to investigate the potential distinct influence of respiratory phases on the modulation of cardiac interoception in individuals with mental or somatic disorders. Numerous studies have linked HEP amplitude to various clinical conditions. For instance, HEP activity has been positively associated with social anxiety (Judah et al., 2018) and anorexia nervosa (Lutz et al., 2019), and negatively associated with depression (Terhaar et al., 2012), post-traumatic stress disorder (Schmitz et al., 2021), depersonalization disorder (Schulz et al., 2015), borderline personality disorder (Schmitz et al., 2020), and chronic pain (Solcà et al., 2020). However, it remains unclear whether these findings reflect an overall variation in HEP activity or a specific change limited to a particular respiratory phase. An interesting exception can be found in the study conducted by Baumert et al. (Baumert et al., 2015), who reported a distinct decrease in HEP amplitude during exhalation compared to inhalation in children with sleep-disordered breathing during sleep. Notably, following the therapy (adenotonsillectomy), the observed effect was a specific increase in HEP amplitude during exhalation, with no changes in HEPs recorded during inhalation. This finding aligns with our current interpretation, suggesting that the effects of an intervention on cardiac interoception are more easily observed during exhalation. Therefore, the potential observation that cardiac interoception is specifically impaired during exhalation in various disorders may facilitate the development of tailored interoceptive interventions. Mindfulness-based practices (Farb et al., 2015; Gibson, 2019; Nord & Garfinkel, 2022; Weng et al., 2021) and slow-breathing techniques (Zaccaro et al., 2018; Zaccaro, Piarulli, et al., 2022) could be designed to specifically target this respiratory phase, aiming to fine-tune cardiac interoception in individual with such disorders.

In conclusion, our data suggest that respiratory phases play a crucial role in influencing cardiac interoception. However, in order to extend this effect to interoception in general and to further explore the hypothesis of an adaptive embodied mechanism that aligns attention with respiration, it is crucial to investigate the impact of respiratory phases on other aspects of interoception beyond cardiac sensations. These domains include gastric interoception (Khalsa et al., 2022; Porciello et al., 2018; Rebollo et al., 2018), interoceptive touch (Crucianelli et al., 2018, 2022; Di Lernia, Cipresso, et al., 2018), and pain perception (Arsenault et al., 2013; Iwabe et al., 2014; Martin et al., 2012; Parrotta et al., 2022). Recognizing this is particularly significant in light of recent models that highlight the role of respiratory activity in shaping mental processes and brain physiology (Allen, Varga, et al., 2022; Boyadzhieva & Kayhan, 2021; Braendholt et al., 2023; Corcoran et al., 2023; Tort et al., 2018; Varga & Heck, 2017). Incorporating these interoceptive dimensions is critical for developing a comprehensive understanding of the reciprocal interactions between the body and the brain (Criscuolo et al., 2022).

## 4. Materials and Methods

### 4.1. Ethics statement

The Institutional Review Board of Psychology of the Department of Psychological, Health and Territorial Sciences, “G. d’Annunzio” University of Chieti-Pescara, Italy, approved the present study (Protocol Number 44_26_07_2021_21016). Participants were kept blind to the experimental aims. The study was conducted in conformity with the Italian Association of Psychology and the Declaration of Helsinki and its later amendments. Before participating in the experiment, all subjects read and signed an informed consent.

### 4.2. Data and code availability

The code and source data used to produce statistics and figures are available in an open repository at the following link: https://github.com/azaccaro90/HEP_attention_respiration_HCT. EEG raw data and individual ERP files can be shared by the corresponding author upon reasonable request if data privacy can be guaranteed according to the rules of the European General Data Protection Regulation (EU GDPR 2016).

### 4.3. Participants

Thirty-four (19 females; 1 left-handed; age: 27.48 ± 4.28 years [mean ± SD]) healthy volunteers participated in the study. The eligibility of each volunteer was verified by self-reported criteria, including: 1) no personal and/or family history of psychiatric, neurological, and somatic disorders; 2) no chronic and/or acute conditions involving the respiratory tracts; 3) no use of any substance interacting with the central nervous system in the previous week; 4) no experience in mindfulness-based meditation and/or breath-control practices. All participants were of normal weight and had normal or corrected-to-normal vision. The study sample size was estimated using G*Power 3.1 software (v3.1.9.7; Faul et al., 2007). Considering a repeated measures ANOVA design, we estimated a medium effect size of partial eta squared (*η*2p) of .06, based on a recent meta-analysis on attentional effects on the HEP (Coll et al., 2021), which corresponds to a Cohen’s f of .25. The significance level was set to alpha = .05, and the desired power (1 - beta) at .85. The total estimated sample size was 26. We excluded 2 participants from the testing of HEPs, regardless of the respiratory phase, due to noisy EEG data (N = 32, 17 females; 1 left-handed; age: 27.57 ± 4.33 years [mean ± SD]). Additionally, we excluded 4 more participants from the testing of HEP changes among respiratory phases due to noisy respiratory data (N = 28, 13 females; 1 left-handed; age: 27.88 ± 4.46 years [mean ± SD]).

### 4.4. Experimental procedure

Participants underwent a baseline and four different tasks in the same experimental session. After a resting-state (baseline), they performed two interoceptive tasks and two exteroceptive tasks, requiring them to focus their attention either inside or outside the body, respectively (“Attention Focus” factor). During both the interoceptive and the exteroceptive tasks, participants had to process either cardiac or respiratory signals (“System” factor). The Baseline was always performed first. Participants were seated for 10 minutes in a resting-state condition with eyes open, watching a fixation cross at the centre of a monitor, and letting their mind wander without focusing on anything in particular (Raichle et al., 2001). Participants were also asked to breathe spontaneously.

The interoceptive tasks were a cardiac task and a respiratory task. The cardiac interoceptive task was the Heartbeat Counting Task (HCT, Pollatos et al., 2005; Schandry & Weitkunat, 1986). Participants had to focus on their cardiac activity and sensations, mentally count and report the number of perceived heartbeats while watching a fixation cross on the screen. Participants were explicitly given the following instructions: 1) not to manually check their heartbeat anywhere on the body; 2) to report only heartbeats they actually felt, rather than those they guessed to occur; 3) to breathe spontaneously and to avoid reducing breath-frequency and breath-holding (apnoea) periods. The HCT consisted of three blocks, each one comprising four HCT trials of the following durations: 25, 35, 45, and 100 seconds (Ardizzi & Ferri, 2018; Di Lernia, Serino, et al., 2018; Ueno et al., 2020). Within each block, the presentation order of the above-mentioned trials was randomized. After each trial, participants verbally reported the number of perceived heartbeats, and rated their performance (cardiac interoceptive confidence) on a 10-points Likert scale ranging from 0 (very bad performance) to 9 (excellent performance). No feedback was provided about their performance. Altogether, participants performed 12 HCT trials. The respiratory interoceptive task was the Breath Counting Task (BCT; Herrero et al., 2018). Participants were asked to focus on their respiratory activity and sensations, mentally count and report the number of their respiratory cycles. The BCT consisted of five blocks, each comprising one trial of 120 seconds. Participants were explicitly asked 1) to breathe spontaneously; 2) to avoid reducing breath-frequency and breath-holding (apnoea) periods; 3) to only be aware of their breathing activity without interfering with it. After each trial, participants verbally reported the number of performed respiratory cycles (each one comprising an inhalation and an exhalation), and rate their performance (respiratory interoceptive confidence) on a Likert scale ranging from 0 (very bad) to 9 (excellent). In addition, they were asked to rate their perceived ease in performing the task without excessively modifying the respiratory rate on a Likert scale ranging from 0 (very difficult) to 9 (very easy). Again, no feedback was provided about their performance. The order of the two interoceptive tasks (HCT and BCT), as well as the order of the blocks in each task, was randomized across participants.

The exteroceptive tasks mirrored the interoceptive tasks and included a cardiac task and a respiratory task. In the cardiac exteroceptive task, participants were presented with digitally constructed heartbeat sounds and instructed to mentally count and report the number of heard heartbeats while breathing spontaneously with eyes open (Cardiac-Tone Counting Task, C-TCT, Wiebking et al., 2015). Similar to the HCT, the C-TCT consisted of three blocks of four randomized trials of different durations (i.e., 25, 35, 45, and 100 seconds). In all blocks, heartbeats were presented at an irregular frequency, with an average frequency of 60 bpm. After each trial, participants were instructed to report the number of heard heartbeats and rate their performance (cardiac exteroceptive confidence). In the respiratory exteroceptive task, participants were presented with audio samples of respiratory sounds (Breath-Tone Counting Task, B-TCT; adapted from Herrero et al., 2018). Like the BCT, the B-TCT consisted of five blocks, each one comprising one trial of 120 seconds. Audio samples comprised clearly distinguishable inhalation and exhalation sounds, performed at irregular frequency, with an average rate of 18 breaths per minute. After each trial, participants were asked to report the number of heard respiratory cycles and to rate their performance (respiratory exteroceptive confidence). They also had to rate their perceived ease in performing the task without excessively modifying the respiratory rate, as in the BCT. The order of the two exteroceptive tasks (C-TCT and B-TCT), as well as the order of the blocks in each task, were randomized across participants.

To avoid presentation order effects, the two interoceptive tasks and the two exteroceptive tasks were presented in a counterbalanced order between participants. Each task was preceded by a brief training. Before starting a task, participants had to verbally confirm that they could not feel their heartbeat through the respiratory belt, in case the belt was wrongly mounted too tight (Zaccaro et al., 2022). After each task, to prevent fatigue and maintain focus, participants were allowed to take short breaks. To ensure a similar number of HEP epochs, the baseline and the four tasks (HCT, BCT, C-TCT, and B-TCT) were all of the same duration, lasting 10 minutes each. EEG, ECG, and respiratory signals were simultaneously recorded throughout the entire experiment. A schematic representation of the experimental design is depicted in Figure 5. All tasks were administered using the E-Prime 3.0 software (Psychology Software Tools, Pittsburgh, PA, USA) connected to a TriggerStation^TM^ (BRAINTRENDS LTD 2010, Rome, Italy).

**Figure 5.**
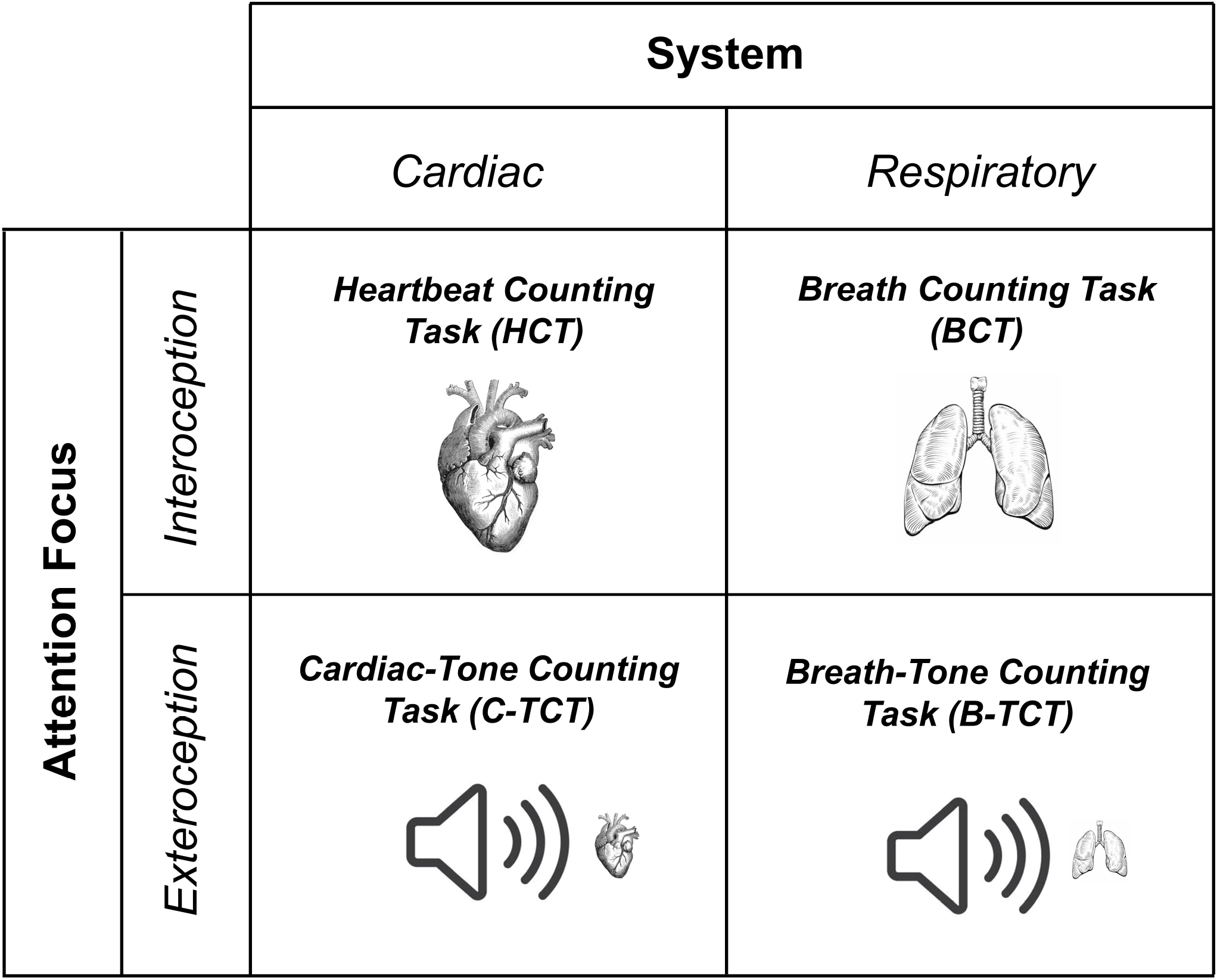
Experimental design. Schematic representation of the experimental design.

### 4.5. Electrophysiological recording

EEG signals were acquired using a 64-channel BrainAmp EEG system (BrainCap MR, BrainVision LLC, Garner, NC, USA). Electrodes were positioned on the scalp following the International 10-20 system. Electrode impedance was kept below 10 kΩ for all channels. ECG data were recorded from three ECG electrodes: two placed over the left costal margin and the right clavicle, respectively, and the ground located on the right costal margin (MP160, BIOPAC Systems Inc, Goleta, CA, USA). Three additional electrodes, serving as the backup ECG, were integrated in the EEG recording system via an external Electrode Input Box (Brain Products GmbH, BrainVision LLC, Garner, NC, USA). These backup electrodes were placed over the left and right clavicles, with the ground was located on the left costal margin. Breathing activity was recorded via a respiratory belt placed around the chest (respiratory transducer TSD201, BIOPAC Systems Inc, Goleta, CA, USA). The sampling rate was 2.5 kHz for all signals. Band-pass filtering was applied from .016 to 250 Hz, along with 50-Hz notch filtering. Subsequently, all signals were down-sampled to 512 Hz.

### 4.6. ECG data analysis

ECG data were processed using custom MATLAB code (R2021a, MathWorks Inc, Natick, MA, USA). The ECG signal was high-pass filtered at .1 Hz to remove baseline fluctuations. R-peaks were detected in the ECG trace with the Pan-Tompkins algorithm (Pan and Tompkins, 1985; Sedghamiz, 2014). To measure HRV, the time series of the distance between consecutive R-peaks (tachogram) were computed, and ectopic beats were corrected using a point process model (Citi et al., 2012). This procedure corrected very few heartbeats (baseline: range 0-4; HCT: 0-16; BCT: 0-9; C-TCT: 0-11; B-TCT: 0-9). Kubios software (Kubios HRV Standard, v3.5; Tarvainen et al., 2014) was used to process the tachogram and extract average Heart Rate (HR) and relevant frequency-domain Heart Rate Variability (HRV) features of interest: log power in the High Frequency band (HFlog - .15-.4 Hz) (Lewis et al., 2012; Malik et al., 1996), Heart Rate Variability (HRV) total power, and Low Frequency/High Frequency ratio (LF/HF) (Supplementary Material 1).

### 4.7. Respiratory data analysis

Respiratory data were processed using custom MATLAB scripts (R2021a, MathWorks Inc, Natick, MA, USA). The respiratory signal was first high-pass filtered at .1 Hz to remove baseline wander. Respiratory phases were detected following a validated procedure (Grund et al., 2022; Power et al., 2020). Firstly, outliers in the respiratory time series (i.e., values exceeding three scaled median absolute deviations from the local median within a moving 1-second window) were linearly interpolated with their neighbouring values. The data were then smoothed with a 1-second-window filter (Savitzky and Golay, 1964), and z-scored. Inhalation and exhalation onsets were identified using the MATLAB function *findpeaks* as the local minima (through) and the local maxima (peaks), respectively. Peaks and troughs were required to be at least 2 seconds apart, and the minimum prominence of the interquartile range of the z-scored data was multiplied by a value ranging from .3 to .8, depending on the participant’s respiratory trace (Grund et al., 2022). A respiratory cycle was defined as the interval from one inhalation onset (referred to as “through”) and the subsequent through. Respiratory cycles with outlier durations (i.e., different from two times the median) were excluded. The number of rejected respiratory cycles was as follows: 6 ± 5 [mean ± SD] for the baseline, 9 ± 6 for the HCT, 4 ± 3 for the BCT, 12 ± 6 for the C-TCT, and 8 ± 7 for the B-TCT. The total number of retained artifact-free respiratory cycles for the entire sample was 3997 for the baseline (143 ± 37 [mean ± SD]), 3969 for the HCT (142 ± 31), 4053 for the BCT (145 ± 34), 4590 for the C-TCT (164 ± 28), and 4440 for the B-TCT (159 ± 27). Inhale and exhale onsets and offsets were visually inspected, and a set of respiratory features of interest (Zaccaro et al., 2022) were calculated, including breathing rate, average inhale duration, average exhale duration, and Inhalation/Exhalation (I/E) ratio (Supplementary Material 1).

### 4.8. EEG data analysis

The EEG data were pre-processed using the EEGLAB toolbox (v2022.1; Delorme & Makeig, 2004), and custom MATLAB code (R2021a, MathWorks Inc, Natick, MA, USA). Initially, the EEG data were filtered using a Hamming windowed FIR filter in the frequency range of .5 - 45 Hz (Al et al., 2020, 2021). Then, a visual inspection was performed to manually remove the local artefacts and noisy channels, and segments with poor signal quality were manually rejected. Independent Component Analysis (ICA) was applied to the retained signal to visualize and manually remove sources of heartbeat, ocular, and muscle artefacts (FastICA; Hyvärinen, 1999). Physiological channels (ECG and respiratory channel) were excluded from ICA decomposition, as were previously identified noisy channels. The number of noisy channels was minimal, typically less than 5, and these channels were always peripheral, known not to contribute to HEP generation (Coll et al., 2021). The Independent Components (ICs) were visually inspected, and three main indicators were used for rejection: 1) the IC activation time series; 2) the IC map over the scalp; and 3) the IC power spectrum (Delorme & Makeig, 2004). Particular attention was given to the Cardiac Field Artifact (CFA), as it highly impacts HEP activity (Coll et al., 2021). ICs contaminated by the CFA were identified by plotting the R-peak events over the IC activations time series. If an IC’s its activation was time-locked to the ECG R-peak, it was removed (Al et al., 2020, 2021; Zaccaro et al., 2022) (Fig. S8). To facilitate this visual inspection, the cross-correlation between each IC time course and the ECG channel was calculated (MATLAB function *corrcoef*). The number of removed ICs ranged from 2 to 15 (7 ± 3 [mean ± SD]), while the number of CFA-related components ranged from 0 to 3 (1 ± 1 [mean ± SD]). After ICA, the previously identified noisy EEG channels were interpolated using their neighbouring channels (Junghöfer et al., 2000). The pruned EEG signals were then re-referenced to the average reference (Zaccaro et al., 2022), as recommended when investigating brain-heart interplay (Candia-Rivera et al., 2021).

HEPs were computed from EEG signals time-locked to the R-peak of the ECG using the ERPLAB toolbox (v9.00; Lopez-Calderon & Luck, 2014). ECG R-peak latencies were identified with the HEPLAB toolbox (Perakakis, 2019). Following R-peaks detection, manual correction of mis-detected peaks was performed (Al et al., 2021; Zaccaro et al., 2022). EEG data were epoched between −100 and 600 msec, with the R-peak event as the zero-point temporal reference (Coll et al., 2021). Epochs were not baseline-corrected (Petzschner et al., 2019). This decision was made to exclude confounding residual CFA evoked by ECG waves that precede the R-peak (P and Q-waves). In addition, late components of HEPs have been reported up to 620 ms after the R-peak, and avoiding baseline correction prevents the removal of those late effects of interest evoked by the previous heartbeat (Petzschner et al., 2019; Schulz et al., 2013, 2015). Epochs followed by an R-peak by less than 600 ms were excluded to avoid overlap between the HEP activity and the following R-peak residual CFA (Babo-Rebelo et al., 2019; Zaccaro et al., 2022). The number of epochs excluded for this reason was generally low (20 ± 60 [mean ± SD]), except for one participant with heart rate higher than 90 bpm. Epochs were also excluded if they exceeded a peak-to-peak threshold of 100 *μ*V (Blankenship et al., 2018; Villena-González et al., 2017; Zaccaro et al., 2022), using a moving window threshold function (window size: 200 ms; overlap: 50%) (Lopez-Calderon & Luck, 2014). The number of rejected epochs exceeding this threshold was very low (4 ± 10 [mean ± SD] for the baseline, 4 ± 11 for the HCT, 2 ± 6 for the BCT, 3 ± 9 for the C-TCT, and 3 ± 10 for the B-TCT). To test the hypothesis that HEP activity is specifically modulated during the HCT, compared to the other tasks, retained epochs were separately averaged within participants for each task, resulting in four HEP bins named as follows: cardiac interoception from the HCT, cardiac exteroception from the C-TCT, respiratory interoception from the BCT, and respiratory exteroception from the B-TCT. To further investigate whether hypothesis that specific modulations of HEP activity during the HCT are related to the respiratory phases, HEP epochs were assigned to inhalation and exhalation, based on which respiratory phase their respective R-peaks occurred. Then, HEPs corresponding to inhalation and exhalation were computed for each participant during the baseline and each task, by averaging epochs assigned to the respective respiratory phase. This resulted in ten HEP bins named as follows: baseline inhale and baseline exhale for the baseline; cardiac interoception inhale and cardiac interoception exhale for the HCT; cardiac exteroception inhale and cardiac exteroception exhale for the C-TCT; respiratory interoception inhale and respiratory interoception exhale for the BCT; respiratory exteroception inhale and respiratory exteroception exhale for the B-TCT.

### 4.9. Task accuracy assessment

Task accuracy was calculated following validated procedures (Candia-Rivera et al., 2022; Garfinkel et al., 2015; Hart et al., 2013; Schandry, 1981). HCT accuracy was computed based on the number of counted heartbeats in relation to the real number of heartbeats recorded with the ECG during each trial. The formula was:

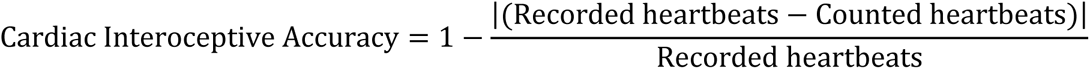

Interoceptive accuracy scores range from 0 to 1, where higher scores indicate better performance. The accuracy of each trial was then averaged over the 12 HCT trials (Candia-Rivera et al., 2022; Garfinkel et al., 2015; Hart et al., 2013). Similarly, task accuracy for the C-TCT was calculated as follows:

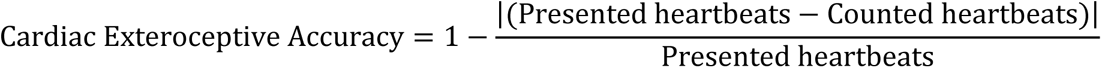

based on the number of counted heartbeats with respect to the number of presented heartbeats sounds within each trial. Similarly, task accuracy for the BCT and the B-TCT were calculated based on the number of the counted respiratory cycles with respect to the number of breaths detected with the respiratory belt and the number of presented respiratory sounds, respectively. Related formulas were:

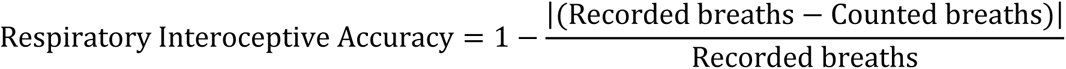

and

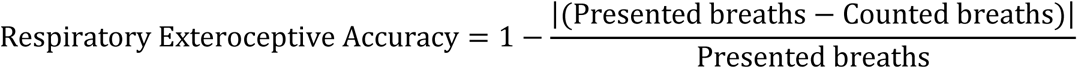

The computed accuracy scores in each trial were then averaged over the 5 respiratory trials. The accuracy score ranges between 0 and 1, where higher scores indicate better performance. For each of the four tasks, self-reported subjective performance (confidence) was averaged among all trials relevant for the respective condition.

### 4.10. Psychometric questionnaires

The Five Facet Mindfulness Questionnaire (FFMQ, Baer et al., 2006; Italian version by Giovannini et al., 2014), and the Multidimensional Assessment of Interoceptive Awareness (MAIA, Mehling et al., 2012; Italian version by Calì et al., 2015) were administered via the Qualtrics online platform (Qualtrics, Provo, UT, USA). In the FFMQ, participants were asked to answer to 39 statements on a 5-point Likert scale ranging from 1 (never) to 5 (always). The FFMQ questionnaire is composed of five subscales and a total dispositional (trait) mindfulness score. It measures the tendency to stay present with perceptions, thoughts, emotions, and actions without distraction, to be non-judgmental of one’s own experience, and to perceive emotions without reacting. In the present study, we focused on the FFMQ total score to correlate HEP amplitude with trait mindfulness (Verdonk et al., 2021). In the MAIA, participants were asked to answer to 37 statements on a 6-point Likert scale ranging from 0 (never) to 5 (always). The MAIA questionnaire consists of eight scales, each assessing different facets of interoception. Given previous association with HEP activity (Baranauskas et al., 2017; Billeci et al., 2021), in the present study we focused on the Trusting and Not-Worrying scales of the MAIA. The trusting scale assesses the degree to which participants experience their body as safe and trustworthy, while the tendency not to worry or experience emotional distress with bodily sensations of pain and/or discomfort.

### 4.11. Statistical analyses

HEP differences between tasks were statistically assessed with the Factorial Mass Univariate ERP Toolbox (FMUT; Fields, 2017; Fields & Kuperberg, 2020) in the MATLAB environment (R2021a, MathWorks Inc, Natick, MA, USA). The FMUT extends the Mass Univariate ERP Toolbox (MUT; Groppe et al., 2011a, 2011b), and applies ERP mass univariate approaches and cluster-based corrections to factorial ANOVAs. To test our first hypothesis of HEP activity modulations induced by the focus of attention, we performed a 2×2 repeated measures ANOVA with the mean HEP amplitude as dependent variable. The within-participants factors were System (cardiac vs. respiratory) and Attention Focus (interoceptive vs. exteroceptive). Using a permutation-based approach (*FclustGND* function), we sought significant main effects of both System and Attention Focus, as well as a System by Attention Focus interaction effect on HEP amplitude. The permutation-based cluster mass F-test involves testing clusters for significance, instead of individual electrodes or time points. Firstly, spatially (i.e., neighbouring channels) and temporally (i.e., contiguous time-points) adjacent points with a p-value below .05 are included into a cluster. The cluster mass statistic for each cluster is then calculated by summing all the F-values within that specific cluster. The null distribution for the cluster mass statistic is estimated through a Monte Carlo permutation procedure, in which condition labels are randomly shuffled a large number of times, and the largest cluster-level statistic obtained at each randomization is entered into the null distribution. Finally, statistical significance is determined by comparing the experimentally observed cluster-level statistics with the randomly-generated null distribution. Observed clusters exceeding the 1 - alpha percentile of the distribution (two-tails) are considered significant (Fields, 2017; Fields & Kuperberg, 2020). To perform permutation tests in factorial ANOVA designs, the FMUT uses the permutation of residuals method, which is associated to reduced Type I error rate (Anderson & Ter Braak, 2003; Still & White, 1981; Winkler et al., 2014). For instance, to test for the main effect of Attention Focus (interoceptive vs. exteroceptive) in a within-subjects design, the FMUT first calculates the average of both levels of System (cardiac and respiratory) for each participant. Then, the FMUT performs a one-way ANOVA between the average values of the interoceptive and exteroceptive Attention Focus. To test the 2×2 interaction (System by Attention Focus), FMUT first calculates the difference between the two levels of Attention Focus for each level of System (cardiac interoception minus cardiac exteroception, and respiratory interoception minus respiratory exteroception) for each subject. Then, it conducts a one-way ANOVA comparing these difference values across the two levels of System (cardiac differences vs. respiratory differences). This approach provides correction for multiple comparisons in time and space and offers high statistical power for effects that are broadly distributed across the scalp and time (Groppe et al., 2011b). Planned t-tests on mean HEP amplitude were conducted using paired cluster-based permutation t-tests implemented in the MUT between each pair of interests (Groppe et al., 2011a). Specifically, we planned to compare HCT vs. C-TCT, HCT vs. BCT, BCT vs. B-TCT, and C-TCT vs. B-TCT. As the HEP waveform clearly showed two distinct trends, an early negative deflection between 150 and 349 msec after the R-peak, and one late more positive oscillation from around 350 to 600 msec (Fig. S9), we selected two time-windows of interest covering the HEP epoch for statistical analyses (Baranauskas et al., 2017; Kumral et al., 2022; Schulz et al., 2018; Simor et al., 2021). The early time-window (i.e., 150-349 msec after the R-peak) corresponds to the cardiac systole, which is known to be highly affected by residual CFA (Coll et al., 2021). Conversely, the late time-window (i.e., 350-600 msec after the R-peak) corresponds to the cardiac diastole, when the CFA is at a minimum (Babo-Rebelo et al., 2019; Babo-Rebelo, Richter, et al., 2016; Dirlich et al., 1997). To reduce the number of statistical comparisons and increase power, we conducted tests on the mean HEP amplitude separately for the early (i.e., 150-349 msec) and the late (i.e., 350-600 msec) time-windows. For the analysis, we excluded peripheral and posterior electrodes that were more susceptible to noise contamination and known to contribute less to HEP modulations related to interoceptive attention (Coll et al., 2021). Consequently, in line with recent works (Coll et al., 2021; Fittipaldi et al., 2020), we focused on HEP differences over 16 channels in a fronto-central region of interest (Fpz, Fp1, Fp2, AFz, AF3, AF4, Fz, F1, F2, F3, F4, FCz, FC1, FC2, FC3, and FC4) (Fig. S10). The number of permutations for each statistical comparison was 10000. The maximum distance value for two electrodes to be included into a cluster was 5.44 cm, assuming a 56 cm circumference head (*chan_hood* = .61).

To test our second hypothesis, which investigates whether HEP activity modulations are related to the respiratory phases, we computed HEP differences between inhalation and exhalation. For this analysis, we conducted a 2×2×2 repeated measures ANOVAs with mean HEP as dependent variable, and System (cardiac vs. respiratory), Attention Focus (interoceptive vs. exteroceptive), and Phase (inhalation vs. exhalation) as within-participants factors. This repeated measures ANOVA was performed over the significant time-window resulting from the testing of the first hypothesis. To further explore respiratory phase-dependent HEP differences across the four tasks, we conducted planned t-tests on mean HEP amplitude using paired cluster-based permutation t-tests implemented in the MUT between each pair of interests (Groppe et al., 2011a). Specifically, we planned to perform the following comparisons: HCT inhale vs. HCT exhale; BCT inhale vs. BCT exhale; C-TCT inhale vs. C-TCT exhale; B-TCT inhale vs. B-TCT exhale; HCT exhale vs. C-TCT exhale; HCT exhale vs. BCT exhale; HCT inhale vs. C-TCT inhale; and HCT inhale vs. BCT inhale.

### 4.12. Respiratory cycle analyses

Circular statistics (Pewsey et al., 2013) enable the assessment of HEPs recorded across the entire respiratory cycle, without differentiating between inhalation and exhalation, while also correcting for variations in the duration of respiratory cycles. Following the circular analysis approach, we examined HEP activity over the entire respiratory cycle, spanning from one inhalation to the subsequent one. To achieve this, we firstly calculated the relative position of the R-peak onset within the respiratory cycle using the following formula, adapted from Al and colleagues (Al et al., 2020):

Rpeak relative position

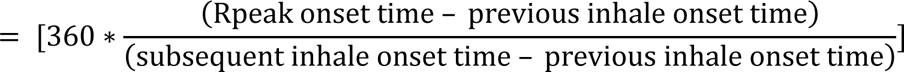

which resulted in values ranging between 0° and 360°, with 0° indicating the onset of inhalation. Subsequently, single-trial angular values were converted to radians (range 0 - 2π), and the corresponding HEP values were derived from electrodes and time-window showing significant changes across the respiratory cycle in the binary analysis. Outliers exceeding three standard deviations from the mean were removed from the dataset. Using a median split method, HEP values were categorized into two bins: high-amplitude HEPs and low-amplitude HEPs, for each participant (Al et al., 2020, 2021). For each participant, the mean angle direction was calculated for both the low-amplitude and high-amplitude HEPs. At the group level, we tested whether the distribution of high and low HEPs deviated from uniform distribution across the respiratory cycle using the Hodges-Ajne test (or Omnibus test, see Park et al., 2020). This test assesses the uniformity of phase distribution of circular data without making assumptions on the distribution of the data (Ajne, 1968). The Hodges-Ajne test yields in the M statistic, which represents the minimum number of data points that can be observed in half of the circle. If the test statistic is smaller than the expected number, the null hypothesis of a uniform distribution is rejected (Ajne, 1968), indicating a non-uniform distribution of data around the respiratory cycle. Next, to assess whether the strength of the respiration-locking significantly differed between high- and low-amplitude HEPs, we applied the Watson-Williams test (Stephens, 1969; Watson & Williams, 1956). This test evaluates whether the mean directions of the two groups are identical or not. Although the Watson-Williams test assumes underlying von Mises distributions with equal concentration parameter, it has demonstrated robustness against deviations from these assumptions (Zar 1999; Berens, 2009). For all circular statistics analyses, we used MATLAB functions implemented in the Circular Statistics Toolbox (Berens, 2009).

To assess respiratory phase-related effects on HEP amplitude, we computed HEPs amplitude based on six equally sized respiration phase bins spanning the 0°-360° interval (0°-60°, 60°-120°, 120°-180°, 180°-240°, 240°-300°, and 300°-360°). Subsequently, a one-way repeated measures ANOVA was conducted to assess significant changes in HEP amplitude across the respiratory cycle. Planned t-tests comparisons were performed between inhalation phase bins (0°-60°, 60°-120°, 120°-180°) and exhalation phase bins (180°-240°, 240°-300°, 300°-360°) (number of t-tests: 9).

Throughout the manuscript, p-values were adjusted for multiple testing, when appropriate, using the Benjamini and Yekutieli procedure (FDR, Benjamini and Yekutieli, 2001) with an FDR threshold set at p = .05. Effect sizes were estimated by computing Cohen’s d and partial eta squared (*η*2p) indices. Statistical analyses of behavioural and physiological data were conducted using repeated measures ANOVAs. Correlations between behavioural and psychometric data with HEP values were performed using Pearson’s correlation. All statistical analyses were conducted in jamovi (v2.3.21; The jamovi project, 2022).

## CRediT authorship contribution statement

**Andrea Zaccaro:** Conceptualization, Methodology, Software, Validation, Formal analysis, Investigation, Data curation, Visualization, Writing-Original Draft, Writing-Review & Editing. **Francesca della Penna**: Investigation, Formal analysis, Data curation, Writing-Review & Editing. **Elena Mussini**: Software, Data Curation, Writing-Review & Editing. **Eleonora Parrotta**: Software, Data Curation, Writing-Review & Editing. **Mauro Gianni Perrucci**: Methodology, Software, Data curation, Resources, Writing-Review & Editing. **Marcello Costantini**: Visualization, Supervision, Funding acquisition, Writing-Review & Editing. **Francesca Ferri**: Conceptualization, Methodology, Resources, Supervision, Project administration, Funding acquisition, Visualization, Writing-Review & Editing.

