## Supplementary_Material for "The interplay between focus of attention, respiratory phases, and the Heartbeat Evoked Potential"

**Supplementary Material 1. Cardio-respiratory features**

**Heart Rate (HR)** (beats/min): the average number of heartbeats in one minute.

**Heart Rate Variability (HRV) total power** (ms2/Hz): the sum of the power values in three frequency bands: High Frequency (HF: .15 - .4 Hz), Low Frequency (LF: .04 - .15 Hz), and Very Low Frequency (VLF: 0 - .04 Hz). It provides an estimation of HRV in the frequency domain.

**High Frequency logarithmic power (HFlog)** (log ms2/Hz): log-transformed power of the HF band (.15-.4 Hz). It serves as a marker of parasympathetic activation, and is a primary indicator of Respiratory Sinus Arrhythmia (i.e., the influences of respiratory cycle on HR).

**Low Frequency/High Frequency ratio (LF/HF)**: the ratio between Low Frequency and High Frequency band. It estimates the balance of sympatho-vagal systems. A decrease in this ratio indicates a prevalence of parasympathetic activity.

**Breathing rate** (breaths/min): the average number of breathing cycles in one minute.

**Inhale duration** (sec): the average duration of the inhalation phases of respiration.

**Exhale duration** (sec): the average duration of the exhalation phases of respiration.

**Inhalation/Exhalation (I/E) ratio**: the ratio between the average inhale duration and the average exhale duration.

**Supplementary Material 2. Cardio-respiratory control analyses**

To investigate the possible explanatory role of cardiac and respiratory physiology on HEP modulations, we performed a series of additional analyses. We tested for differences in the mean ECG amplitude across the four tasks with a 2x2 repeated measures ANOVA within the time-window of significant HEP differences (i.e., 350-600 msec after the R-peak). We found no main effect for either System or Attention Focus on the ECG amplitude (main effect System: F_1,31_ = .022, p = .884, 𝜂2p = .001; main effect Attention Focus: F_1,31_ = .82, p = .372, 𝜂2p = .003). In addition, we found no interaction effect between System and Attention Focus (F_1,31_ = 2.626, p = .115, 𝜂2p = .008). This null result revealed that the ECG amplitude was unchanged across the four tasks (Fig. S2). As an additional control, we correlated the mean ECG amplitude recorded during the HCT with the mean HEP amplitude observed during the HCT over significant electrodes and time-window for the HCT vs. C-TCT comparison. We found no correlation between ECG amplitude and HEP activity during the HCT (Pearson’s r = .11, p = .55), in line with previous research indicating that these variables are unrelated when the CFA is adequately removed from the EEG data (Buot et al., 2021). We then applied 2x2 repeated measures ANOVA to compare participants’ HR across the four tasks. We found a significant main effect of System (F_1,31_ = 6.6, p = .015, 𝜂2p = .175), and a significant System by Attention Focus interaction effect (F_1,31_ = 38.31, p < .001, 𝜂2p = .553). No main effect of Attention Focus was found (F_1,31_ = 2.28, p = .142, 𝜂2p = .068). Planned t-tests revealed reduced HR for the HCT, compared to both the C-TCT (t_31_ = -3.68, p < .001, Cohen’s d = .65), and the BCT (t_31_ = -5.38, p < .001, Cohen’s d = .951), as well as reduced HR during the C-TCT than during the B-TCT (t_31_ = 2.28, p = .03, Cohen’s d = .402). No changes were detected between the BCT and the B-TCT (t_31_ = 1.04, p = .304, Cohen’s d = .185) (Fig. S3A). This result is in line with a recent work by Candia-Rivera and colleagues (2022), suggesting that reduced HR while focusing on heartbeats may be part of the physiological mechanisms related to cardiac interoceptive attention. When focusing on HFlog power, we found a significant main effect of Attention Focus (F_1,31_ = 14.905, p < .001, 𝜂2p = .325), but no significant main effect of System (F_1,31_ = .318, p = .577, 𝜂2p = .010), and no interaction effect between System and Attention Focus (F_1,31_ = 3.038, p = .091, 𝜂2p = .089). Thus, HFlog power was higher during both the interoceptive tasks (HCT and BCT) compared to the exteroceptive tasks (C-TCT and B-TCT) (Fig. S3B). The same pattern emerged for the HRV total power, showing a significant main effect of Attention Focus (F_1,31_ = 12.872, p = .001, 𝜂2p = .293), but not main effect of System (F_1,31_ = .051, p = .824, 𝜂2p = .002), and no significant System by Attention Focus interaction (F_1,31_ = 3.132, p = .087, 𝜂2p = .092) (Fig. S3C). When testing LF/HF, no significant main effect of System (F_1,31_ = .181, p = .673, 𝜂2p = .006) or Attention focus (F_1,31_ = .004, p = .953, 𝜂2p = < .001) was found, and no interaction effect was observed (F_1,31_ = .075, p = .786, 𝜂2p = .002). Moving to the breath frequency, only a significant main effect of Attention Focus was detected (Main effect Attention Focus: F_1, 31_ = 61.399, p < .001, 𝜂2p = .664; Main effect System: F_1,31_ = 2.226, p = .146, 𝜂2p = .067; System by Attention Focus: F_1,31_ = .374, p = .545, 𝜂2p = .012). This indicates reduced breath frequency during both the interoceptive tasks, compared to the exteroceptive tasks (Fig. S3D). The same pattern was found examining the average inhale duration (Main effect Attention Focus: F_1,31_ = 27.007, p < .001, 𝜂2p = .466; Main effect System: F_1,31_ = 2.587, p = .118, 𝜂2p = .077; System by Attention Focus: F_1,31_ = .007, p = .933, 𝜂2p < .001, Fig. S3E), the average exhale duration (Main effect Attention Focus: F_1,31_ = 20.982, p < .001, 𝜂2p = .404; Main effect System: F_1,31_ = 1.338, p = .256, 𝜂2p = .041; System by Attention Focus: F_1,31_ = .055, p = .816, 𝜂2p = .002, Fig. S3F), and the I/E ratio (Main effect Attention Focus: F_1,31_ = 4.732, p < .037, 𝜂2p = .132; Main effect System: F_1,31_ = 1.35, p = .254, 𝜂2p = .042; System by Attention Focus: F_1,31_ = .136, p = .714, 𝜂2p = .004, Fig. S3G).

To assess if HEP activity during the performance of the HCT was influenced by cardiac and respiratory physiology, we performed a series of correlation analyses testing for relationships between HEP and different cardio-respiratory features of interest. Specifically, we correlated the mean HEP amplitude with mean HR, HFlog power, HRV total power, and LF/HF ratio as cardiac features, and breathing rate, average inhale duration, average exhale duration, and I/E ratio as respiratory features. None of these features significantly correlated with HEP amplitude (Pearson’s correlation HR, r = .013, p = .943; Pearson’s correlation HFlog, r = .099, p = .591; Pearson’s correlation HRV total power, r = .175, p = .337; Pearson’s correlation LF/HF, r = -.272, p = .133; Pearson’s correlation breathing rate, r = .335, p = .061; Pearson’s correlation average inhale duration, r = -.316, p = .078; Pearson’s correlation average exhale duration, r = -.136, p = .458; Pearson’s correlation I/E ratio, r = -.206, p = .258).

**Supplementary Material 3. Single-trial cardio-respiratory analyses**

We controlled for HR influences on HEPs across the four tasks by assessing single-trial level data through linear mixed-effects model analyses on 91980 trials. These trials were clustered around 32 participants. This mixed-effect model allowed us to test for the effects of System and Attention Focus factors on single-trial HEP activity while simultaneously controlling for instantaneous HR, resulting in a significant gain in statistical power, as previously reported (Kliegl et al., 2010; Ambrosini et al., 2019; Zaccaro et al., 2022). To calculate instantaneous HR (bpm) for each heartbeat, we based it on the inter-beat interval from the current R-peak to the following R-peak. For each heartbeat, single-trial HEP activity was calculated by averaging the voltage of the significant channels in the time-window of significant observed HEP effects. The mixed-effects model was implemented with jamovi (v2.3.21; The jamovi project, 2022) via the General Analyses for the Linear Model (GAMLj) module (Gallucci, 2019). We used the Residual Maximum Likelihood (REML) method to fit the model. The Akaike Information Criterion (AIC) was utilized to include or exclude factors from the model (Portet, 2020), and the best fitting model was determined by the AIC value being at least 2 points lower than the previous model (Burnham and Anderson, 2002). To facilitate model convergence, we centred the covariance scaling of the variables (Ambrosini et al., 2019; Zaccaro et al., 2022). Regarding continuous variables, outliers were excluded from the final model, specifically values lying outside three standard deviations from the mean. We assessed the statistical significance of predictors using t-tests with Satterthwaite’s approximation of degrees of freedom. For each parameter included in the best fitting model, we reported the estimated coefficient (b), standard error (SE), t-statistic, and p-value. To compare the four tasks, we computed 14 models, starting from the simplest model and then adding factors stepwise (Table S4). The best fitting model, Model 10, included a fixed intercept among participants (ID), the parameter for the interaction between Attention Focus (interoception vs. exteroception) and System (cardiac vs. respiratory), and the Intercept and Attention Focus factors as random effects. Notably, including the System factor as random effect led to model overfitting. The R-notation formula of the best fitting model was:

$$HEP \sim1 + System*Attention Focus + \left( 1+Attention Focus \right| ID)$$

As for the main analysis, the final model confirmed the interaction between System and Attention Focus (b = -.071, SE = .026, t = -2.722, p = .006). We also found no main effect of System (b = -.003, SE = .013, t = -.251, p = .901), Attention Focus (b = .059, SE = .034, t = 1.727, p = .094), or the intercept (b = -.014, SE = .026, t = -.542, p = .592) on HEP activity. This result indicates a non-significant effect of the HR on HEP activity across the four tasks (Fig. S4).

**Supplementary Material 4. Cardio-respiratory control analyses across the respiratory cycle**

We performed additional analyses to control for the potential confounding influence of cardio-respiratory physiology on HEP activity and to support the assumption that observed differences in HEP amplitude between inhalation and exhalation were not driven by heart stroke volume modifications between respiratory phases, as indexed with ECG signal amplitude and HR (Buot et al., 2021). We first tested mean ECG amplitude changes across tasks and respiratory phases with a 2x2x2 repeated measures ANOVA within the time-window of significant HEP differences (i.e., 350-600 msec after the R-peak). We found a significant main effect of Phase on ECG amplitude (F_1,27_ = 15.84, p < .001, 𝜂2p = .37), but no main effect of Attention Focus (F_1,27_ = .093, p = .762, 𝜂2p = .003), and System (F_1,27_ = <.001, p = .986, 𝜂2p = <.001). Moreover, no interactions were found to be significant (System by Attention Focus: F_1,27_ = .825, p = .372, 𝜂2p = .03, Attention Focus by Phase: F_1,27_ = .269, p = .608, 𝜂2p = .01; System by Phase: F_1,27_ = .429, p = .518, 𝜂2p = .016; System by Attention Focus by Phase: F_1,27_ = 1.667, p = .207, 𝜂2p = .058). Overall, consistent with previous findings (Zaccaro et al., 2022), ECG signal amplitude was higher during the exhalation phase, compared to inhalation (Fig. S5, Fig. S6A). Similarly, we tested for HR changes across System, Attention Focus, and Phase with a 2x2x2 repeated measures ANOVA. The analysis revealed a significant main effect of System (F_1,27_ = 5.4, p = .028, 𝜂2p = .167), but no main effect of Attention Focus (F_1,27_ = 1.817, p = .189, 𝜂2p = .063), and Phase (F_1,27_ = .013, p = .91, 𝜂2p = <.001). Furthermore, no interactions were observed between Attention Focus and Phase (F_1,27_ = .499, p = .486, 𝜂2p = .018) and System and Phase (F_1,27_ = 1.995, p = .169, 𝜂2p = .069). However, the System by Attention Focus interaction was significant (F_1,2_ = 31.264, p < .001, 𝜂2p = .537), as well as the three-way System by Attention Focus by Phase interaction (F_1,27_ = 4.821, p < .037, 𝜂2p = .151). Planned t-tests revealed that HR at exhalation during the HCT was significantly lower than HR recorded at exhalation during the C-TCT (t_27_ = -3.247, p = .003, Cohen’s d = .614), and significantly lower than HR at exhalation during the BCT (t_27_ = -5.359, p < .001, Cohen’s d = 1.01). To support the conclusion that observed HEP changes across phases were not related to concurrent changes in HR, planned t-tests showed no significant differences in HR between inhale and exhale during the HCT (t_27_ = .55, p = .587, Cohen’s d = .104). However, in contrast to changes in HEP activity, HR at inhalation during the HCT differed significantly from HR at inhalation during the C-TCT (t_27_ = -3.24, p = .003, Cohen’s d = .612) and from the HR at inhalation during the BCT (t_27_ = -3.9, p < .001, Cohen’s d = .736) (Fig. S6B). These results suggest that HR changes across the four tasks were not driven by a specific phase of the respiratory cycle, as observed for the HEP changes. Nevertheless, these additional findings made it necessary to control for the instantaneous HR effects over HEP amplitude among respiratory phases at the single-trial level (Supplementary Material 5).

To assess if observed HEP changes across respiratory phases during the HCT were driven by changes in cardiac and respiratory physiology, we performed a series of correlation analyses testing for relationships between observed mean HEP differences among respiratory phases (ΔHEP: HEP exhalation minus HEP inhalation computed over significant electrodes and time-window) and cardio-respiratory features of interest. Specifically, we correlated ΔHEP with mean HR, ΔHR (HR exhalation minus HR inhalation), ΔECG (ECG exhalation minus ECG inhalation), HFlog power, HRV total power, and LF/HF ratio as cardiac features, and breathing rate, average inhale duration, average exhale duration, and I/E ratio as respiratory features. However, none of these cardio-respiratory features showed a significant relationship with the observed ΔHEP effects (Pearson’s correlation HR, r = -.177, p = .366; Pearson’s correlation ΔHR, r = .127, p = .518; Pearson’s correlation ΔECG, r = .211, p = .282; Pearson’s correlation HFlog, r = -.024, p = .904; Pearson’s correlation HRV total power, r = -.044, p = .825; Pearson’s correlation LF/HF, r = -.054, p = .784; Pearson’s correlation breathing rate, r = -.111, p = .576; Pearson’s correlation average inhale duration, r = .134, p = .496; Pearson’s correlation average exhale duration, r = -.009, p = .964; Pearson’s correlation I/E ratio, r = .267, p = .169).

**Supplementary Material 5. Single-trial cardio-respiratory analyses across the respiratory cycle**

We controlled for cardiac influences on HEPs across the respiratory cycle (inhalation vs. exhalation) at the single-trial level during the performance of the HCT. We ran a linear mixed-effects model analysis comprising 17196 trials. Trials were clustered around 28 participants. This mixed-effect model allowed us to test for the effects of respiratory phases on single-trial HEP activity by controlling at the same time for instantaneous HR and trial-based ECG amplitude. Instantaneous HR (bpm) was calculated for each heartbeat based on the inter-beat interval from the current R-peak to the following R-peak. Trial-based ECG amplitude was calculated for each heartbeat by averaging the ECG channel voltage in the significant time window of observed HEP effects (i.e., 350-600 msec after the R-peak). Single-trial HEP activity was calculated for each heartbeat by averaging the voltage of the significant channels in the time-window of significant observed HEP effects. The mixed-effects model was implemented as in Supplementary Material 3. We computed 13 models regressing HEP outcome, starting from the simplest model, and then adding factors stepwise (Table S5). The best fitting model was Model 7, which included a fixed intercept among participants (ID), the parameter for the main effect of Phase (inhale vs. exhale), and the parameter for the main effect of the ECG amplitude. It also included the Intercept and the Phase factors as random effects. The R-notation formula of the best fitting model was:

$$HEP \sim1 + Phase+ECG+ \left( 1 + Phase \right| ID)$$

The final model showed a significant main effect of Phase (b = -.105, SE = .051, t = -2.063, p = .048) and ECG amplitude (b = .487, SE = .154, t = 3.159, p = .002), and no main effect of the intercept (b = .026, SE = .047, t = .562, p = .579) over HEP activity. Notably, no interaction effects were found between HR, ECG amplitude, and respiratory phases, indicating that these covariates do not mediate the effect of the respiratory phase on HEP amplitude during the HCT (Fig. S7).

**Supplementary Material 6. Task accuracy and correlation with the HEP**

Task accuracy was influenced by System and Attention Focus (Table S6). A 2x2 repeated measures ANOVA revealed a significant main effect of System (F_1,31_ = 73.1, p < .001, 𝜂2p = .702), with lower task accuracy in the cardiac condition than the respiratory condition. The main effect of Attention Focus was also significant (F_1,31_ = 108.5, p < .001, 𝜂2p = .778), with lower accuracy in the interoception condition than the exteroception condition. Furthermore, there was a significant System*Attention Focus interaction (F_1,31_ = 110.7, p < .001, 𝜂2p = .781). Planned t-tests revealed lower task accuracy in the HCT compared to both the C-TCT (t_31_ = -10.78, p < .001, Cohen’s d = 1.91) and the BCT (t_31_ = -9.66, p < .001, Cohen’s d = 1.71), as well as lower task accuracy in the B-TCT compared to the C-TCT (t_31_ = -2.6, p = .048, Cohen’s d = .365). No differences were observed between the BCT and the B-TCT (t_31_ = -1.81, p = .08, Cohen’s d = .32). In line with previous findings (Candia-Rivera et al., 2022), HCT accuracy strongly correlated with cardiac interoceptive confidence, as measured by confidence ratings of one's own perceived task performance (Pearson’s r = .706, p < .001), and C-TCT accuracy correlated with cardiac exteroceptive confidence (Pearson’s r = .542, p = .001). In contrast, BCT accuracy did not correlate with respiratory interoceptive confidence (Pearson’s r = .277, p = .125), and B-TCT accuracy did not correlate with respiratory exteroceptive confidence (Pearson’s r = .144, p = .433). This discrepancy is likely due to a systematic underestimation of participants’ ability in the respiratory task accuracy, together with a ceiling effect of high accuracy scores during both the BCT and B-TCT. Regarding the BCT and the B-TCT, participants reported a medium-to-high perceived ease in performing the tasks without significantly altering their respiratory rate (BCT: 7.51 ± .96 [mean ± SD]; B-TCT: 7.04 ± 1.27). We found no relationship between HEP amplitude and cardiac interoceptive accuracy and confidence. Correlation analysis with the overall HEP amplitude showed no significant relationship with cardiac interoceptive accuracy (Pearson’s r = .043, p = .817) or cardiac interoceptive confidence (Pearson’s r = -.006, p = .975). Similarly, HEPs amplitude specifically recorded during inhalation or exhalation did not correlate with cardiac interoceptive accuracy or confidence (HEP inhalation and cardiac interoceptive accuracy: Pearson’s, r = .098, p = .618; HEP inhalation and cardiac interoceptive confidence Pearson’s r = .066, p = .737; HEP exhalation and cardiac interoceptive accuracy Pearson’s r = .05, p = .8; HEP exhalation and cardiac interoceptive confidence Pearson’s, r = -.093, p = .636).

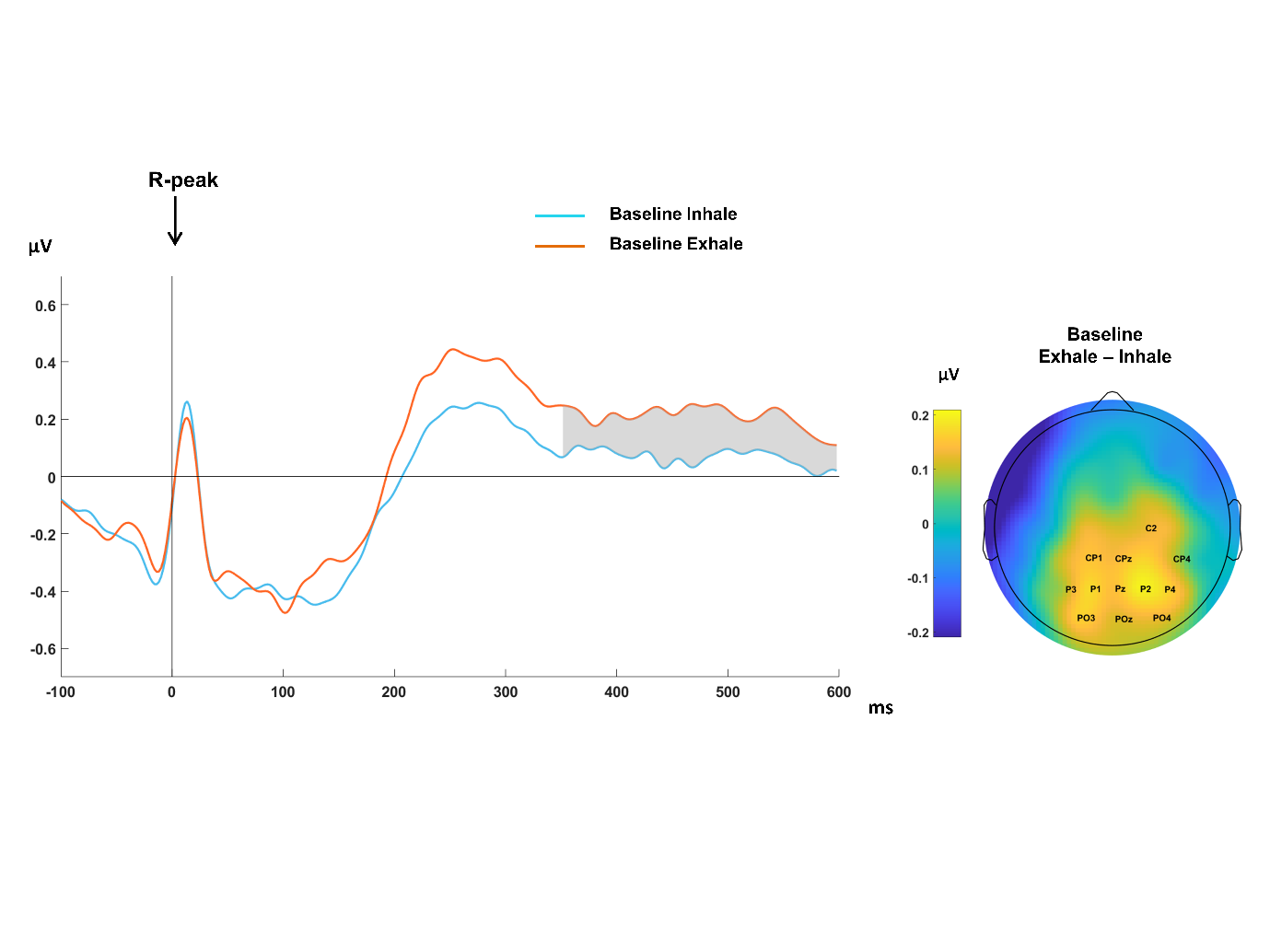

**Supplementary Figure 1. HEP activity is related to the respiratory phase during baseline.** **(left)** Grand-average HEP waveforms pooled for significant electrodes during the baseline. The time courses of the HEP are shown for inhalation (light blue) and exhalation (orange). The grey area marks the time window of significant differences in the contrast between exhalation and inhalation. **(right)** Topographical scalp distribution showing mean HEP differences (350 - 600 msec after the R-peak) between exhalation vs. inhalation during the baseline (significant differences were detected over highlighted electrodes: C2, CP1, CPz, CP4, P3, P1, Pz, P2, P4, PO3, POz, and PO4).

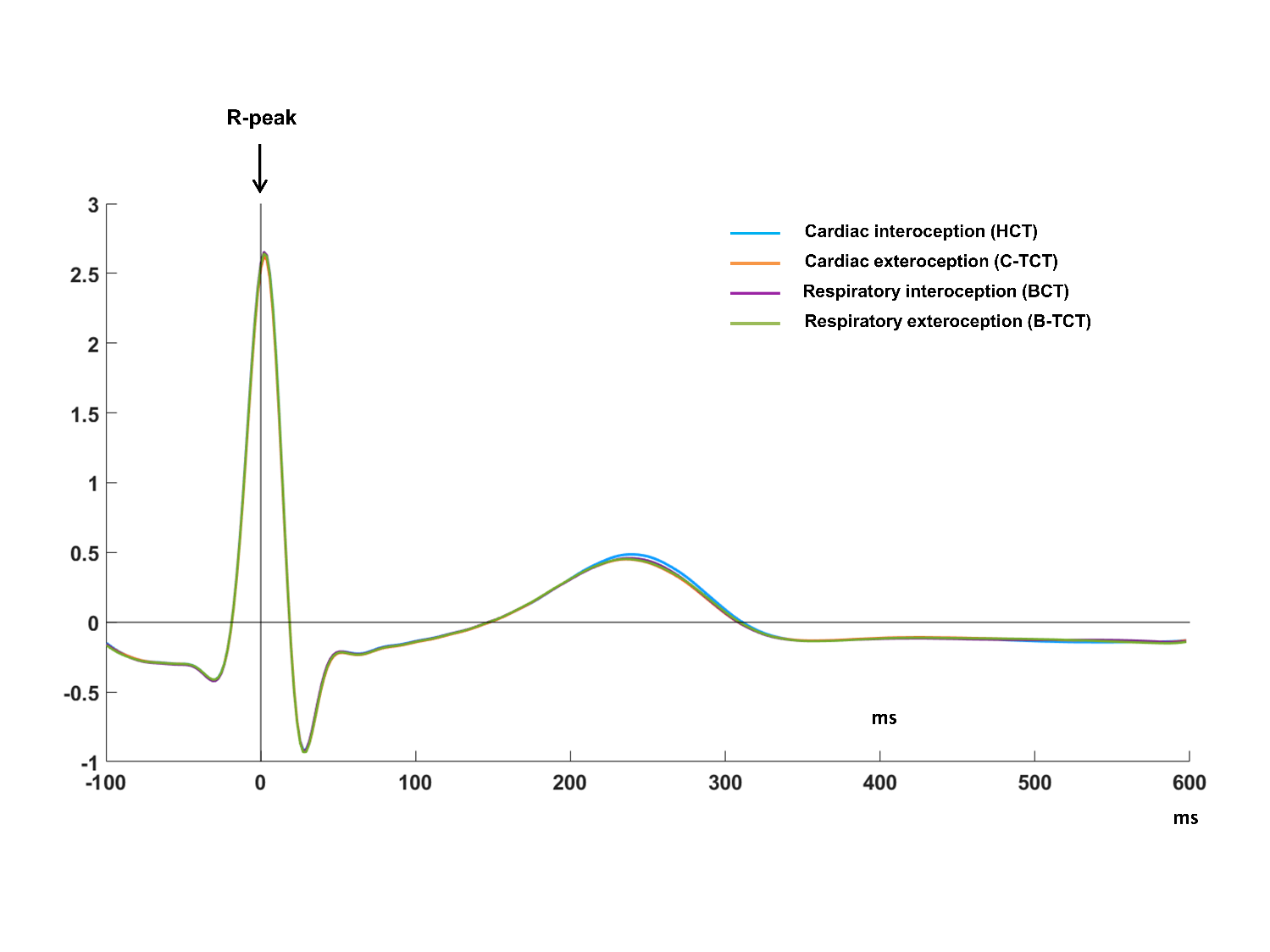

**Supplementary Figure 2. ECG amplitude comparison across System and Attention Focus factors.** Grand average ECG signal time-locked to the R-peak. The time courses of the ECG are shown for the HCT (light blue), C-TCT (orange), BCT (purple), and B-TCT (green). Abbreviations: HCT-Heartbeat Counting Task, C-TCT-Cardiac-Tone Counting Task, BCT-Breath Counting Task, B-TCT-Breath-Tone Counting Task.

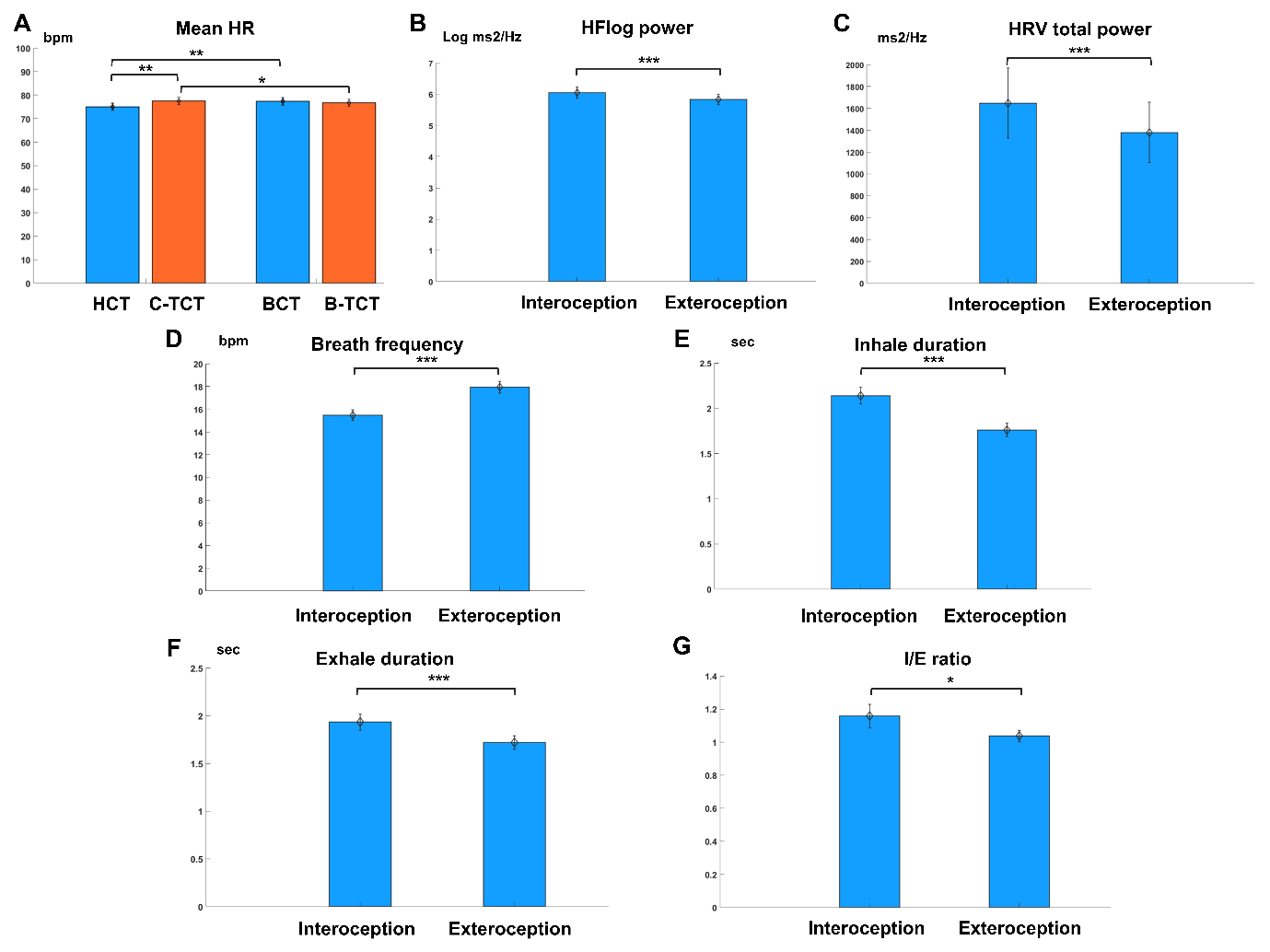

**Supplementary Figure 3. Cardio-respiratory features across the four tasks.** The bar plots display significant effects resulting from the ANOVAs between the Attention Focus and System factors for various cardiac and respiratory parameters. **(A)** Significant planned t-tests elucidating the interaction between Attention Focus and System factors for the mean HR. Significant main effects of the Attention factor are shown in **(B)** for HFlog power, **(C)** for HRV total power, **(D)** for breath frequency, **(E)** for inhale duration, **(F)** for exhale duration, and **(G)** for I/E ratio. Significance is denoted by asterisks (* p < .05; ** p < .01; *** p < .001). Abbreviations: HCT-Heartbeat Counting Task, C-TCT-Cardiac-Tone Counting Task, BCT-Breath Counting Task, B-TCT-Breath-Tone Counting Task, I/E-Inhalation/Exhalation.

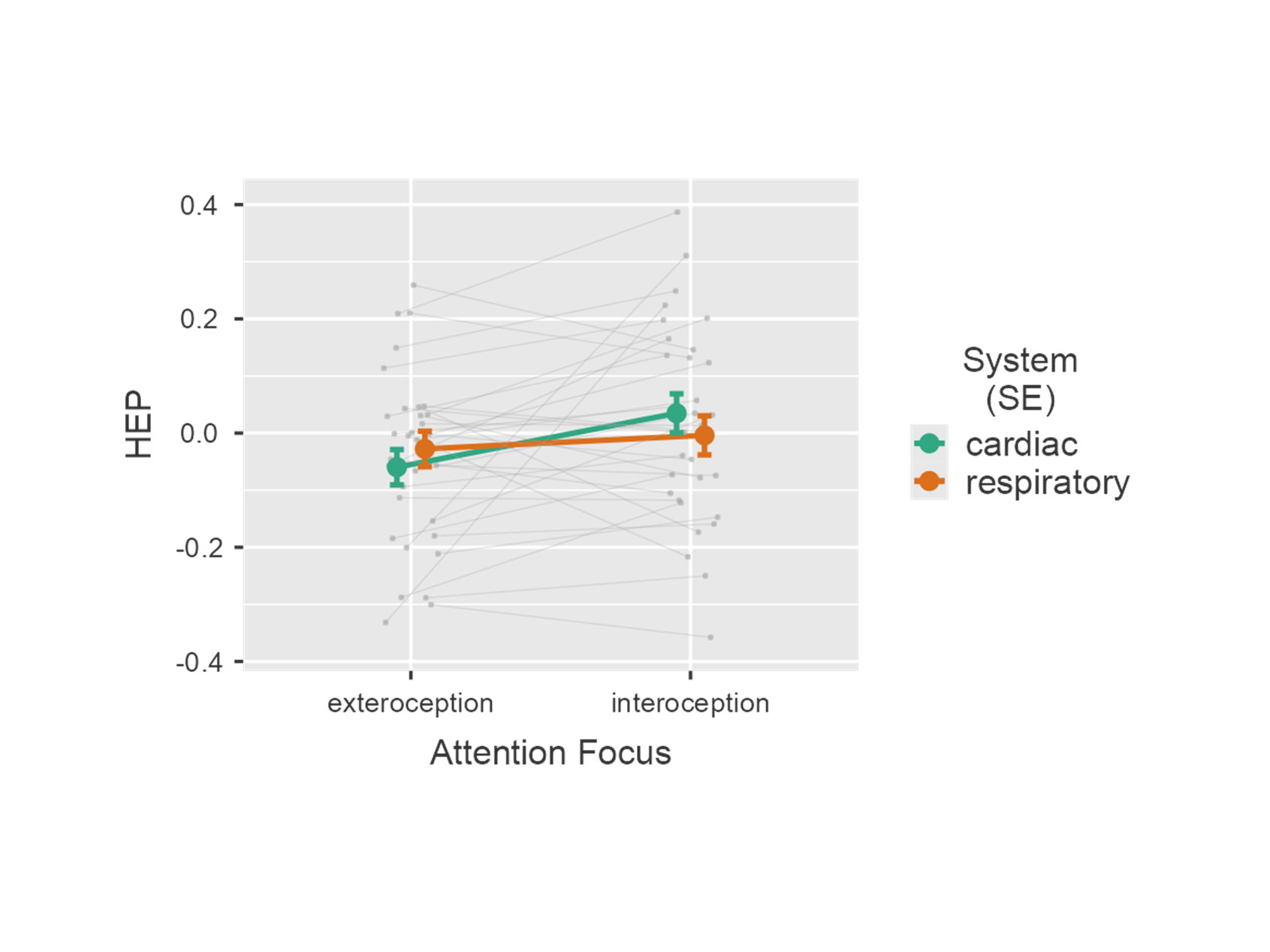

**Supplementary Figure 4. Linear mixed-effects model during the four tasks.** Significant interaction between Attention Focus and System factors on HEP activity across the four tasks. Error bars represent the Standard Error (SE). Random effects are plotted by each participant.

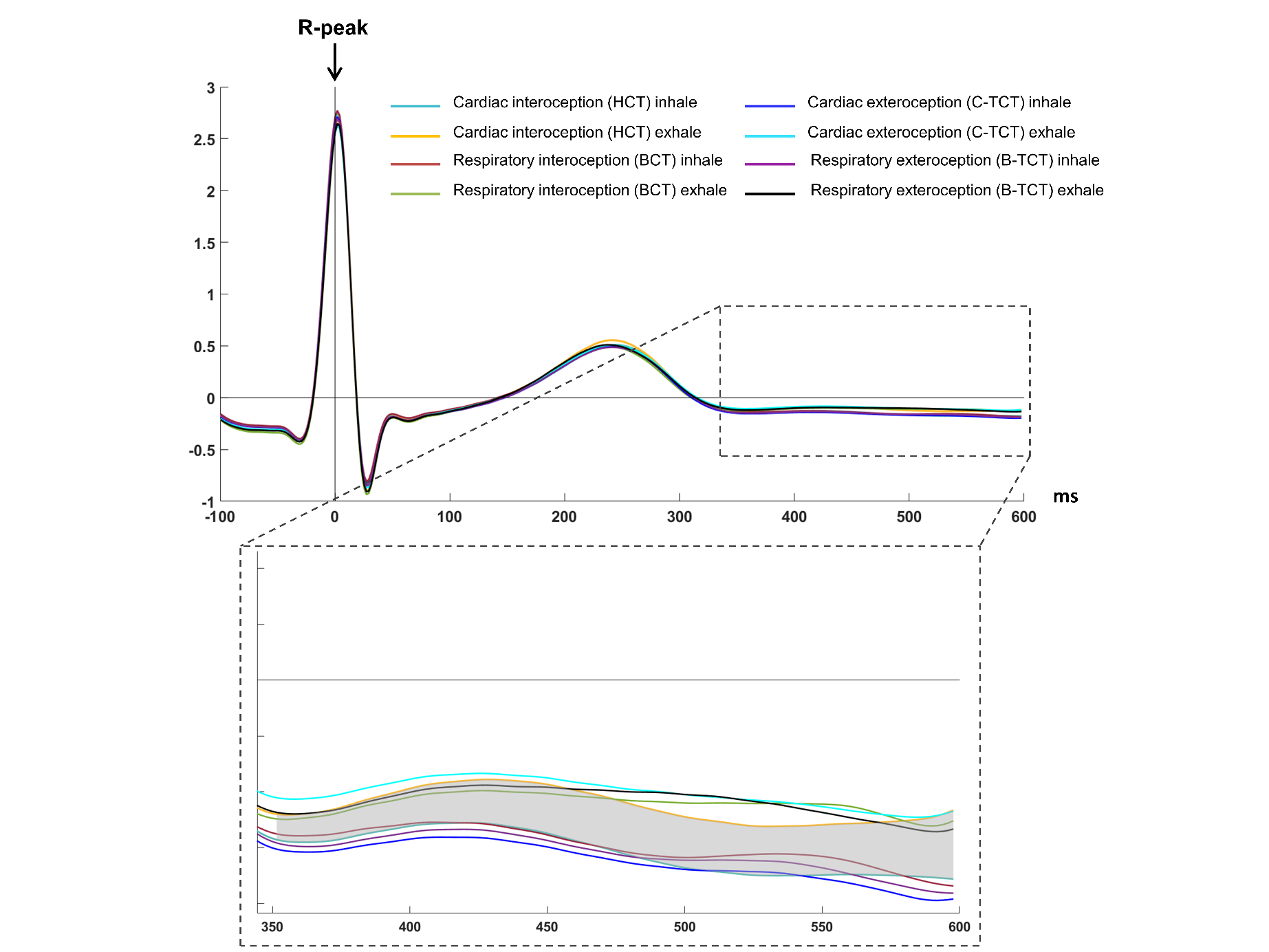

**Supplementary Figure 5. ECG amplitude comparison across System, Attention Focus, and Phase.** Grand average ECG signal time-locked to the R-peak. The time courses of the ECG are shown for the HCT-inhale (turquoise), HCT-exhale (yellow), BCT-inhale (dark red), BCT-exhale (green), C-TCT-inhale (blue), C-TCT-exhale (light blue), B-TCT-inhale (purple), B-TCT-exhale (black). The grey area marks the time window of significant differences in the contrast between the exhale and inhale phase (ΔHEP) during the HCT. Abbreviations: HCT-Heartbeat Counting Task, C-TCT-Cardiac-Tone Counting Task, BCT-Breath Counting Task, B-TCT-Breath-Tone Counting Task.

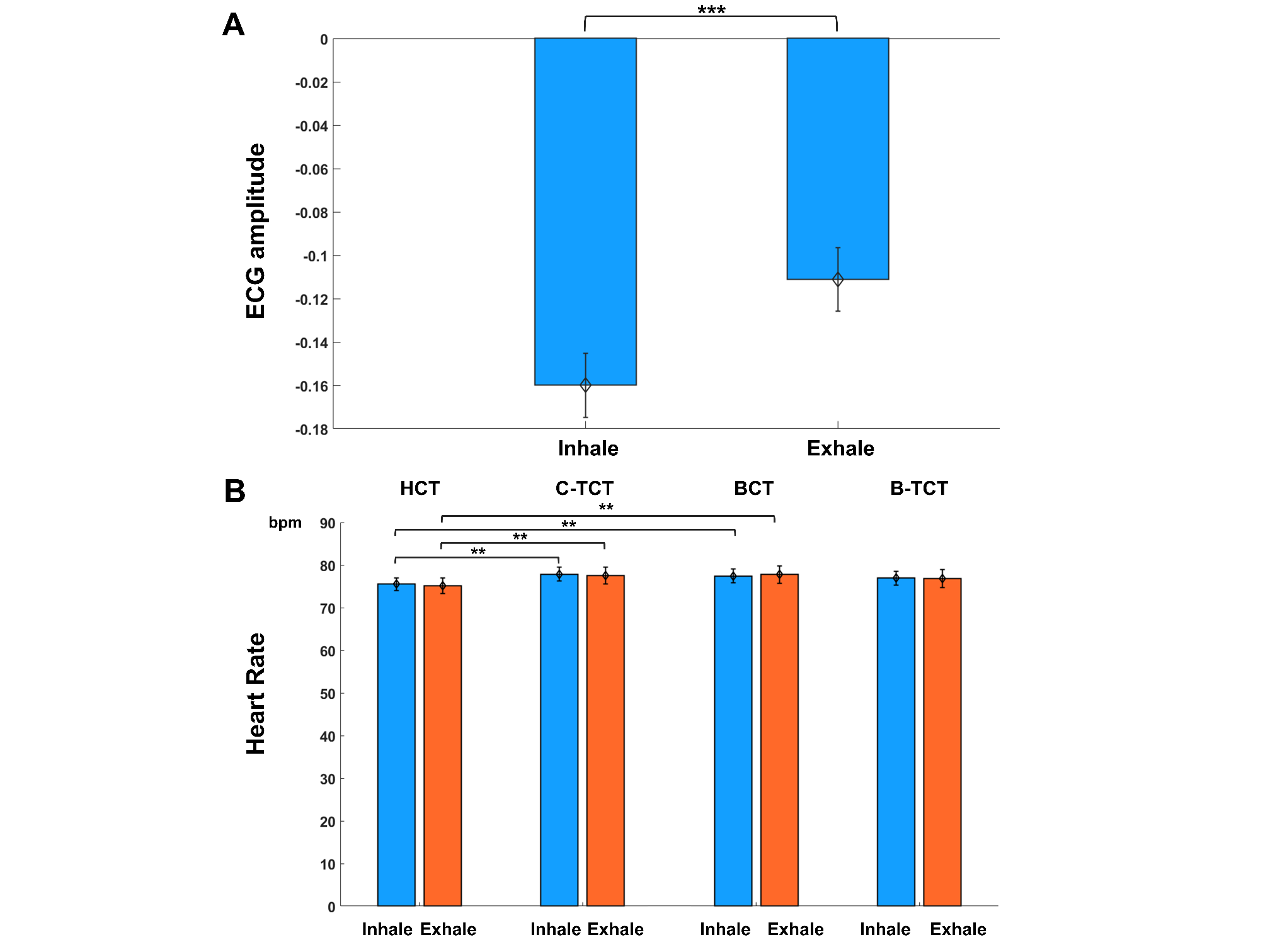

**Supplementary Figure 6. Cardio-respiratory features across the four tasks and respiratory phases.** The bar plots display significant effects resulting from the ANOVAs between the Attention Focus, System, and Phase factors. **(A)** Significant main effect of the Phase factor for the mean ECG amplitude. **(B)** Significant planned t-tests elucidating the triple interaction effect for the mean HR. Significant effects are indicated by asterisks (** p < .01; *** p < .001). Abbreviations: HCT-Heartbeat Counting Task, C-TCT-Cardiac-Tone Counting Task, BCT-Breath Counting Task, B-TCT-Breath-Tone Counting Task.

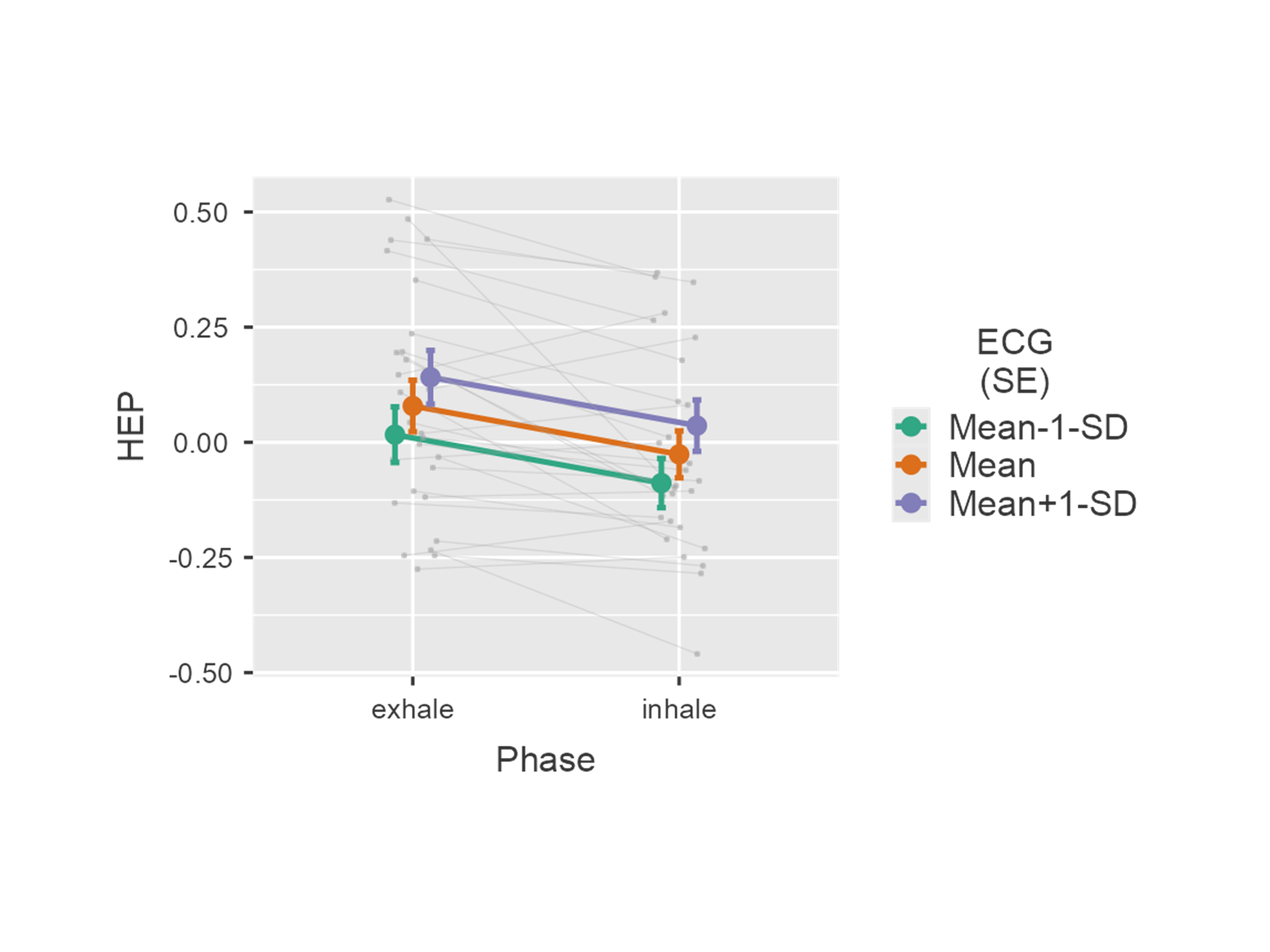

**Supplementary Figure 7. Linear mixed-effects model during the HCT.** Non-significant interaction between Phase and trial-level ECG amplitude on HEP activity during the HCT. Error bars represent the Standard Error (SE). Random effects are plotted by each participant.

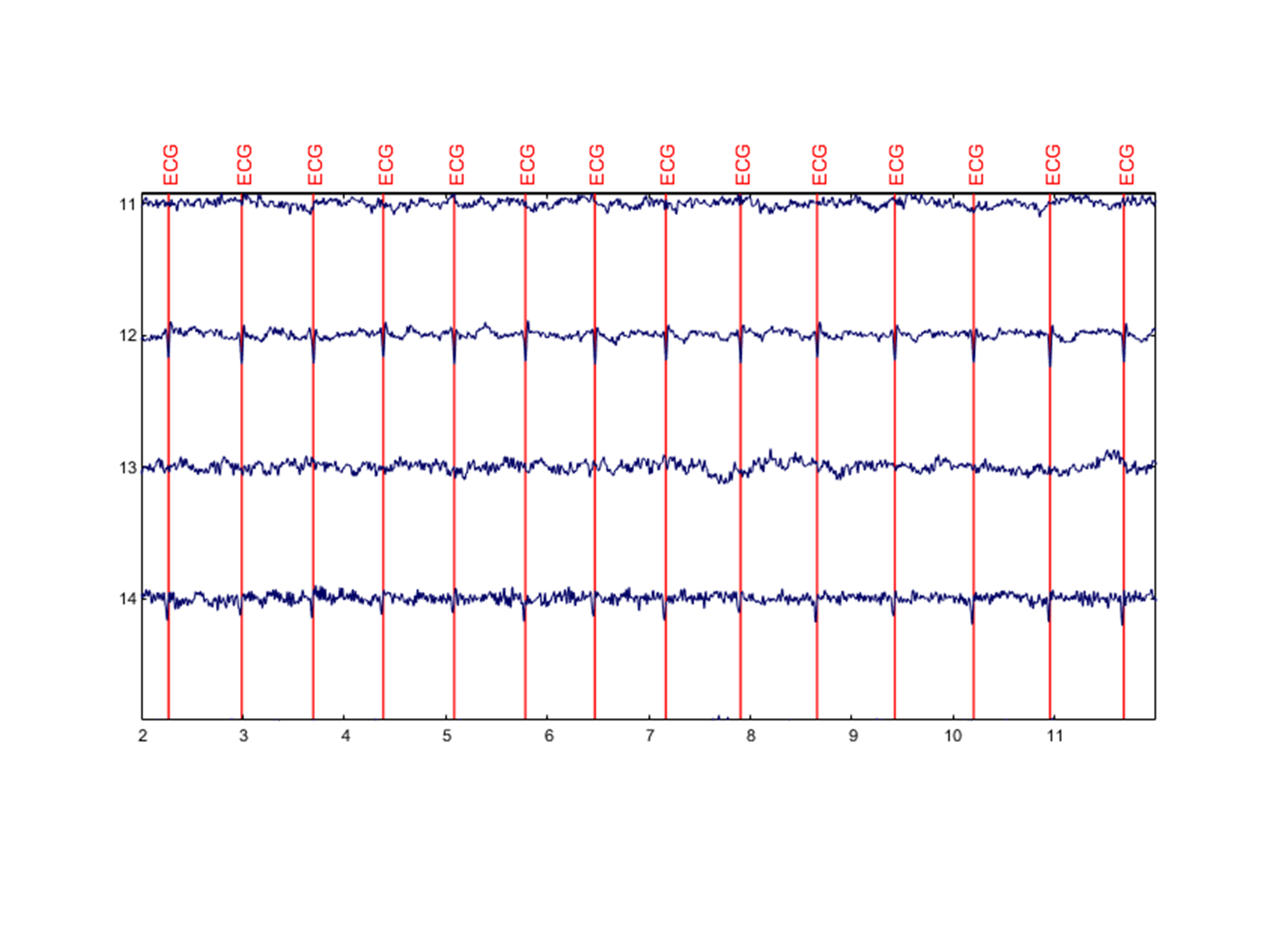

**Supplementary Figure 8. CFA components.** Independent Components (IC) activations time course. The ECG events (red vertical lines) represent the R-peak events. ICs were rejected if their activations were time-locked to the ECG R-peak, hence showing the CFA (IC-12 and IC-14).

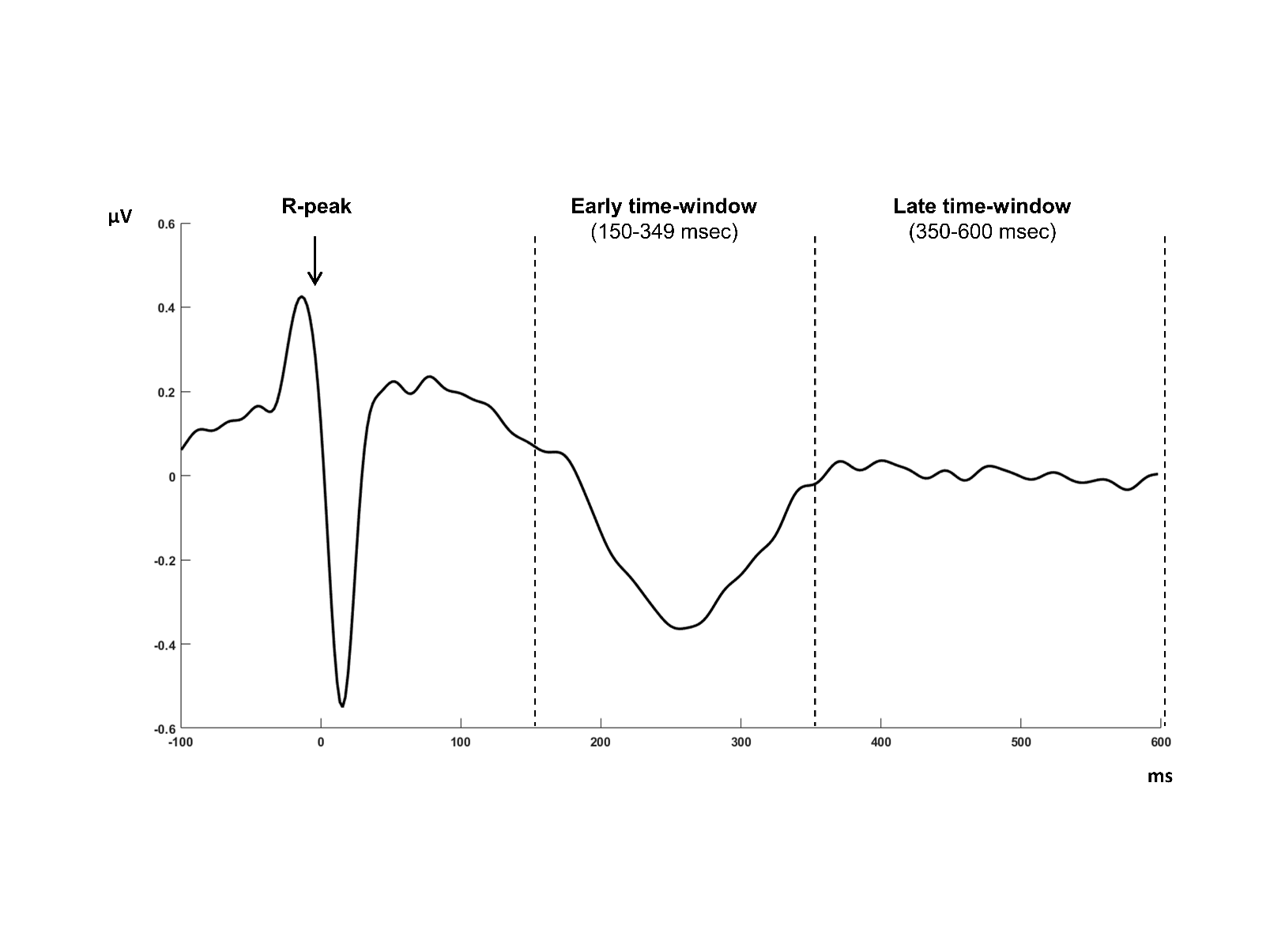

**Supplementary Figure 9. Early and late time-windows of interest.** Grand-average HEP waveforms pooled for ROI electrodes. Time regions of interest for the HEP analyses ranged from 150 to 349 msec (early time-window) and from 350 to 600 msec (late time-window).

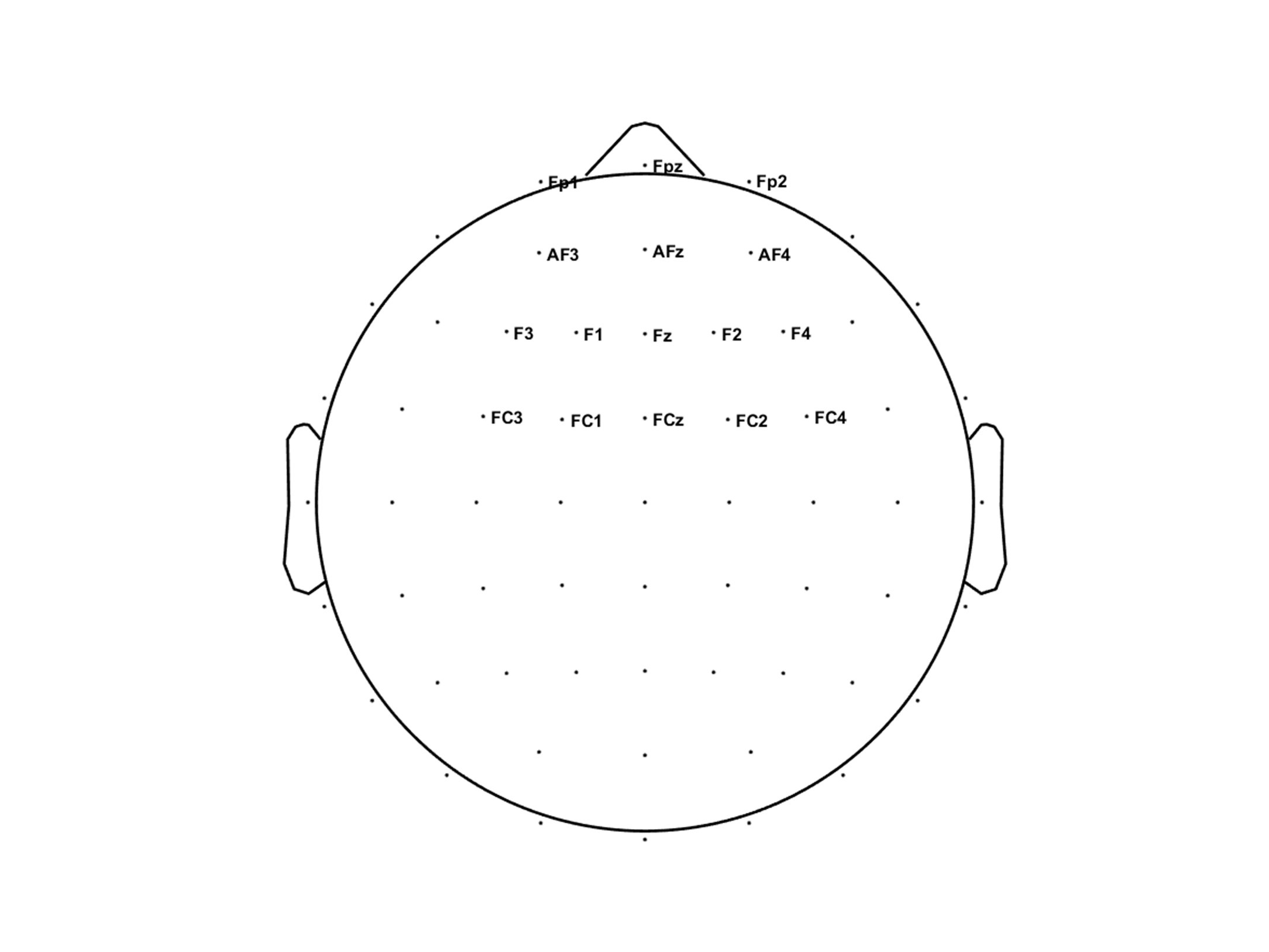

**Supplementary Figure 10. Frontal-central region of interest.** Of the 64 electrodes, 16 located over prefrontal, anterior frontal, frontal, and fronto-central areas were used for the HEP analyses across the four tasks (Fp1, Fpz, Fp2, AF3, AFz, AF4, F3, F1, Fz, F2, F4, FC3, FC1, FCz, FC2, and FC4).

|  | **Interoception** | | | **Exteroception** | | | **Interoception vs. Exteroception** | | |
| --- | --- | --- | --- | --- | --- | --- | --- | --- | --- |
|  | **N** | **M** | **SD** | **N** | **M** | **SD** | **t** | **p** | **Cohen's d** |
| **Cardiac System** | 24831 | 730 | 80 | 25561 | 752 | 79 | 2.299 | .056 | .406 |
| **Respiratory System** | 24552 | 722 | 87 | 24970 | 734 | 84 | .623 | .538 | .11 |

**Supplementary Table 1. HEP epochs across tasks.** Abbreviations: M-Mean, SD-Standard Deviation

|  | **Inhale** | | | **Exhale** | | | **Inhale vs. Exhale** | | |
| --- | --- | --- | --- | --- | --- | --- | --- | --- | --- |
|  | **N** | **M** | **SD** | **N** | **M** | **SD** | **t** | **p** | **Cohen's d** |
| **Baseline** | 10125 | 362 | 74 | 8827 | 315 | 69 | 2.29 | .03 | .432 |
| **HCT** | 8714 | 311 | 74 | 7592 | 271 | 75 | 2.98 | .006 | .563 |
| **BCT** | 10578 | 378 | 62 | 8676 | 310 | 69 | 3.72 | .001 | .703 |
| **C-TCT** | 9264 | 331 | 50 | 8258 | 295 | 54 | 3.25 | .003 | .614 |
| **B-TCT** | 9744 | 348 | 59 | 8374 | 299 | 56 | 3.62 | .001 | .684 |

**Supplementary Table 2. HEP epochs across tasks and respiratory phases.** Abbreviations: HCT-Heartbeat Counting Task, C-TCT-Cardiac-Tone Counting Task, BCT-Breath Counting Task, B-TCT-Breath-Tone Counting Task, M-Mean, SD-Standard Deviation, FDR-False Discovery Rate.

|  | **Interoception** | | | | | | **Exteroception** | | | |
| --- | --- | --- | --- | --- | --- | --- | --- | --- | --- | --- |
|  | **Cardiac Task (HCT)** | | | **Respiratory Task (BCT)** | | | **Cardiac Task (C-TCT)** | | **Respiratory Task (B-TCT)** | |
|  | **M** | | **SD** | | **M** | **SD** | **M** | **SD** | **M** | **SD** |
| **ECG Amplitude** | -.13 | .07 | | -.13 | | .08 | -.13 | .07 | -.13 | .07 |
| **Mean HR** | 75.1 | 8.4 | | 77.42 | | 8.97 | 77.55 | 8.55 | 76.76 | 8.66 |
| **HFlog** | 6.12 | .91 | | 5.99 | | 1.11 | 5.81 | .86 | 5.86 | 1.05 |
| **HRV** | 1736.7 | 1962.1 | | 1562 | | 1772.4 | 1313.7 | 1360.7 | 1446.8 | 1784.1 |
| **LF/HF** | 2.13 | 2.80 | | 2.32 | | 2.79 | 2.19 | 1.67 | 2.23 | 2.08 |
| **Breath Frequency** | 15.73 | 3.19 | | 15.22 | | 3.34 | 18.41 | 3.13 | 17.48 | 3.36 |
| **Inhale Duration** | 2.08 | .63 | | 2.20 | | .67 | 1.70 | .46 | 1.82 | .45 |
| **Exhale Duration** | 1.90 | .47 | | 1.96 | | .61 | 1.68 | .34 | 1.76 | .50 |
| **I/E Ratio** | 1.15 | .48 | | 1.17 | | .38 | 1.01 | .19 | 1.06 | .22 |

**Supplementary Table 3. Cardio-respiratory features across tasks.** Abbreviations: ECG-ElectroCardioGram, HR-Heart Rate, HF-High Frequency, LF-Low Frequency, I/E-Inhalation/Exhalation, HCT-Heartbeat Counting Task, C-TCT-Cardiac-Tone Counting Task, BCT-Breath Counting Task, B-TCT-Breath-Tone Counting Task, M-Mean, SD-Standard Deviation, FDR-False Discovery Rate.

| **N** | **R-notation formula** | **AIC** |
| --- | --- | --- |
| 1 | HEP ∼ 1 + (1\|ID) | 385810 |
| 2 | HEP ∼ 1 + HR + (1\|ID) | 385809 |
| 3 | HEP ∼ 1 + Attention Focus + (1 + Attention Focus\|ID) | 385682 |
| 4 | HEP ∼ 1 + Attention Focus + HR + (1 + Attention Focus\|ID) | 385683 |
| 5 | HEP ∼ 1 + Attention Focus*HR + (1 + Attention Focus\|ID) | 385681 |
| 6 | HEP ∼ 1 + System + (1 + System\|ID) | 385813 |
| 7 | HEP ∼ 1 + System + HR + (1 + System\|ID) | 385813 |
| 8 | HEP ∼ 1 + System*HR + (1 + System\|ID) | 385811 |
| 9 | HEP ∼ 1 + Attention Focus + System + (1 + Attention Focus\|ID) | 385684 |
| **10** | **HEP ∼ 1 + Attention Focus*System + (1 + Attention Focus\|ID)** | **385679** |
| 11 | HEP ∼ 1 + Attention Focus*System + HR + (1 + Attention Focus\|ID) | 385679 |
| 12 | HEP ∼ 1 + Attention Focus*System + HR*Attention Focus + (1 + Attention Focus\|ID) | 385678 |
| 13 | HEP ∼ 1 + Attention Focus*System + HR*System + (1 + Attention Focus\|ID) | 385679 |
| 14 | HEP ∼ 1 + Attention Focus*System + HR*Attention Focus*System + (1 + Phase + System\|ID) | 385679 |

Note: the best fitting model is Model 10, bold

**Supplementary Table 4. Linear mixed-effects models testing between the four tasks.** Abbreviations: ID-Participant, HR-Heart Rate, AIC-Akaike Information Criterion.

| **N** | **R-notation formula** | **AIC** |
| --- | --- | --- |
| 1 | HEP ∼ 1 + (1\|ID) | 74503 |
| 2 | HEP ∼ 1 + HR + (1\|ID) | 74503 |
| 3 | HEP ∼ 1 + ECG + (1\|ID) | 74486 |
| 4 | HEP ∼ 1 + HR + ECG + (1\|ID) | 74486 |
| 5 | HEP ∼ 1 + HR*ECG + (1\|ID) | 74488 |
| 6 | HEP ∼ 1 + Phase + (1 + Phase\|ID) | 74475 |
| **7** | **HEP ∼ 1 + Phase + ECG + (1 + Phase\|ID)** | **74467** |
| 8 | HEP ∼ 1 + Phase*ECG + (1 + Phase\|ID) | 74468 |
| 9 | HEP ∼ 1 + Phase + HR + (1 + Phase\|ID) | 74475 |
| 10 | HEP ∼ 1 + Phase*HR + (1 + Phase\|ID) | 74477 |
| 11 | HEP ∼ 1 + Phase + ECG + HR + (1 + Phase\|ID) | 74467 |
| 12 | HEP ∼ 1 + Phase + ECG*HR + (1 + Phase\|ID) | 74469 |
| 13 | HEP ∼ 1 + Phase*ECG*HR + (1 + Phase\|ID) | 74471 |

Note: the best fitting model is Model 7, bold

**Supplementary Table 5. Linear mixed-effects model during the HCT.** Abbreviations: ID-Participant, HR-Heart Rate, ECG-ElectroCardioGram, AIC-Akaike Information Criterion.

|  | **Interoception** | | | | **Exteroception** | | | |
| --- | --- | --- | --- | --- | --- | --- | --- | --- |
|  | **Cardiac Task (HCT)** | | **Respiratory Task (BCT)** | | **Cardiac**  **Task (C-TCT)** | | **Respiratory Task (B-TCT)** | |
|  | **M** | **SD** | **M** | **SD** | **M** | **SD** | **M** | **SD** |
| **Task Accuracy** | .57 | .22 | .95 | .04 | .98 | .02 | .97 | .04 |
| **Task Confidence** | 5.77 | 1.73 | 7.94 | .74 | 7.93 | .61 | 7.55 | .85 |
| **Respiratory Task Ease** |  |  | 7.51 | .96 |  |  | 7.04 | 1.27 |

**Supplementary Table 6. Task accuracy and confidence.** Abbreviations: HCT-Heartbeat Counting Task, C-TCT-Cardiac-Tone Counting Task, BCT-Breath Counting Task, B-TCT-Breath-Tone Counting Task, M-Mean, SD-Standard Deviation.
